# Immediate Impact of Yogic Breathing on Pulsatile Cerebrospinal Fluid Dynamics

**DOI:** 10.1101/2021.08.09.455068

**Authors:** Selda Yildiz, John Grinstead, Andrea Hildebrand, John Oshinski, William D. Rooney, Miranda M. Lim, Barry Oken

**Author notes:** **Corresponding author.** Selda Yildiz, Department of Neurology, Oregon Health & Science University, 3181 S.W. Sam Jackson Park Road, Portland, OR, 97239, USA.

## Abstract

Cerebrospinal fluid (CSF), a clear fluid bathing the central nervous system (CNS), undergoes pulsatile movements, and plays a critical role for the removal of waste products from the brain including amyloid beta, a protein associated with Alzheimer’s disease. Regulation of CSF dynamics is critical for maintaining CNS health, and increased pulsatile CSF dynamics may alter brain’s waste clearance due to increased mixing and diffusion. As such, understanding the mechanisms driving CSF movement, and interventions that influence its resultant removal of wastes from the brain is of high scientific and clinical impact. Since pulsatile CSF dynamics is sensitive and synchronous to respiratory movements, we are interested in identifying potential integrative therapies such as yogic breathing to regulate and enhance CSF dynamics, which has not been reported before. Here, we investigated the pre-intervention baseline data from our ongoing randomized controlled trial, and examined whether yogic breathing immediately impacts pulsatile CSF dynamics compared to spontaneous breathing. We utilized our previously established non-invasive real-time phase contrast magnetic resonance imaging (RT-PCMRI) approach using a 3T MRI instrument, and computed and rigorously tested differences in CSF velocities (instantaneous, respiratory, cardiac 1^st^ and 2^nd^ harmonics) at the level of foramen magnum during spontaneous versus four yogic breathing patterns. In examinations of 18 healthy participants (eight females, ten males; mean age 34.9 ± 14 (SD) years; age range: 18-61 years), we discovered immediate increase in cranially-directed velocities of instantaneous-CSF 16% - 28% and respiratory-CSF 60% - 118% during yogic versus spontaneous breathing, with most statistically significant changes during deep abdominal breathing (28%, p=0.0008, and 118%, p=0.0001, respectively). Further, cardiac pulsation was the primary source of pulsatile CSF during all breathing conditions except during deep abdominal breathing, when there was a comparable contribution of respiratory and cardiac 1st harmonic power [0.59 ± 0.78], demonstrating respiration can be the primary regulator of CSF depending on individual differences in breath depth and location. Further work is needed to investigate the impact of sustained training yogic breathing on increased pulsatile CSF dynamics and brain waste clearance for CNS health.

## 1. Introduction

### 1.1. Cerebrospinal Fluid

Cerebrospinal fluid (CSF) is one of the two discrete fluid compartments of the brain along with interstitial fluid (ISF), and is crucial for the health of central nervous system (CNS). With the advances in imaging technologies and recent research efforts ^1–11^, it is clear that CSF is more than a mechanical cushion for the CNS and a vehicle for distribution of nutrients and hormones through the CNS. CSF movement ^12–16^ and CSF-ISF exchange ^2,4,17–19^ during wakefulness, sleep and/or anesthesia recently have received particular interest for their implications on pathological states involving CSF. For instance, CSF together with ISF plays an essential role for the removal of solutes and metabolic wastes from the brain interstitium ^1,3,4^ including amyloid beta ^20,21^, a peptide associated with Alzheimer’s disease ^22^, the most common form of dementia contributing to ~60-70% of ~50 million dementia cases worldwide ^23^. Understanding the mechanisms driving CSF movement, and interventions that influence and enhance its resultant removal of waste products from the brain is therefore of high scientific and clinical impact.

CSF movement is driven by pressure changes in CNS vascular system due to cardiac pulsation (~1 Hz) ^2,7,24–32^ and respiration (0.1~0.3 Hz) ^12–16,33–38^, and is influenced by transient effects such as coughing ^14,39–41^, and body posture ^42,43^. A topic of current interest involving CSF dynamics is identifying the primary regulator(s) of CNS fluids or solute movement within subarachnoid spaces, ventricles, deep brain parenchyma ^2,7,12,14,36,37,44^. Recent studies have implicated 1) forced inspiration in humans ^12^, 2) cardiac pulsation with some contribution from respiration in humans ^37^, and 3) cardiac pulsation in rodents^7^, as major drivers of CSF flow. Recent work by extension has also examined the magnitude, direction, and sensitivity of CSF movement to respiratory performances and locations ^14–16,45,46^, and more recently, low-frequency oscillations (e.g., vasomotion; ~ <0.1 Hz) ^9,47,48^ including during sleep ^9^. For instance, in a recent study conducted while subjects sleeping in an MRI scanner, Fultz and colleagues ^9^ demonstrated that CSF flow oscillations during non-rapid eye movement (NREM) sleep were (5.52 dB) larger and slower (0.05 Hz vasomotion) compared to wakefulness (0.25 Hz respiratory), and suggested that increased pulsatile CSF dynamics during sleep may alter the brain’s waste clearance due to increased mixing and diffusion ^2,49^.

In short, CSF movement ^15,16,50^ and removal of solutes ^1,8,18,19,47,51–53^ from the brain is a topic of high clinical impact. Yet, further research is needed for investigating CSF movement under voluntarily controlled conditions to better understand potential therapies for regulating and enhancing CSF dynamics, which has not been investigated. To this end, we designed a study to investigate yogic breathing to modulate and enhance pulsatile CSF dynamics, with the long-term goal of determining whether regular practice of yogic breathing is an effective intervention to aid in the brain’s waste clearance in order to optimize CNS health.

### 1.2. Yogic Breathing

Yogic breathing (*pranayama* ^54^; fourth limb of the -traditional- eight-limb path yoga practice ^54,55^) consists of a variety of breathing techniques performed with conscious control. An appealing modality for healthcare management purposes, yogic breathing is effective for reducing stress and anxiety ^56–58^, lowering blood pressure ^59^, improving asthma conditions ^60^, and improving response to cancer ^61^. A key principle of a regular yogic breathing practice is to make the breath slower, deeper, and rhythmical, which is associated with the self-regulatory mechanism and health-benefits ^57,58,62^. Documented effects ^62^ of slow breathing for instance cover respiratory, cardiovascular, and cardiorespiratory autonomic nervous systems. One commonly studied mechanism ^62^ for the health benefits of yogic breathing is its balancing effect on autonomic nervous system through parasympathetic activation. Since CSF is sensitive to respiratory dynamics ^12–14,37^, and based on our investigations, we believe another potential mechanism for the benefits of yogic breathing is its influence on pulsatile CSF dynamics, which to date has not been reported.

We have recently developed a non-invasive real-time phase-contrast MRI (RT-PCMRI) approach ^14^ that quantifies the influence of both respiration and cardiac pulsations on the (magnitude and direction of) instantaneous CSF velocities in absolute units [cm/s], which provides a unique opportunity to study the impact of yogic breathing practices on CSF dynamics. We have utilized this RT-PCMRI in a recent randomized controlled trial (RCT) that aims to investigate effects of two separate 8-week yogic breathing interventions on pulsatile CSF dynamics. While the RCT aims to investigate long-term impact of yogic breathing, we herein present the pre-intervention baseline data, prior to randomization, to demonstrate the immediate impact of yogic breathing on pulsatile CSF dynamics compared to spontaneous breathing. Briefly, to achieve this, we computed instantaneous-CSF (iCSF) velocities acquired with RT-PCMRI during spontaneous breathing and four yogic breathing practices (for a total of five breathing conditions). We then separated iCSF into three components: respiratory (rCSF), cardiac 1^st^ (c_1_CSF) and 2^nd^ harmonics (c_2_CSF), and rigorously tested the differences between spontaneous versus four yogic breathing conditions.

The goal of this study is then two-fold: (1) to quantify and compare the immediate impact of four yogic breathing practices versus spontaneous breathing on (magnitude and direction of) velocities of iCSF, rCSF, c_1_CSF, and c_2_CSF, and (2) to quantify the relative contribution of rCSF versus c_1_CSF and c_2_CSF during each breathing condition to determine the primary regulator of CSF in all breathing conditions.

## 2. Materials and Methods

### 2.1. Participants

The study was approved by the Institutional Review Board of the Oregon Health & Science University (OHSU), and the full ongoing RCT was registered at the Clinicaltrials.gov (ID # NCT03858309). We received verbal and written informed consent from all study subjects prior to all study procedures. We recruited healthy participants from the Portland metropolitan area using OHSU’s study participation opportunities website, Oregon Center for Clinical and Translational Research Institute (OCTRI) research match for recruitment, flyers throughout the OHSU campus and communities in Portland, and social media (Facebook). We aimed to enroll participants 18 to 65 years of age who were able and available for study activities including undergoing non-invasive MRI scans, had no current or previous regular practice of mind-body therapies focusing on breath awareness and/or training (e.g., yoga, meditation, Tai-Chi, Qi-Gong), and were in good health without any history of neurological disorders, sleep disorders, respiratory disorders, problems with heart, circulatory system, and lungs. See **Table S1** for a full list of RCT inclusion/exclusion criteria. Of the 65 participants contacted for the study, 56 were phone screened, 26 were enrolled, 21 completed the baseline procedures (September-October, 2019 at OHSU), and 18 were included in final baseline data analysis (N=18, eight females, ten males; mean age: 34.9 ± 14 (SD) years; age range: 18-61 years). See **Fig. 1** for the study flow chart, and **Table 1** for the study group characteristics (N=18).

**Figure 1.**
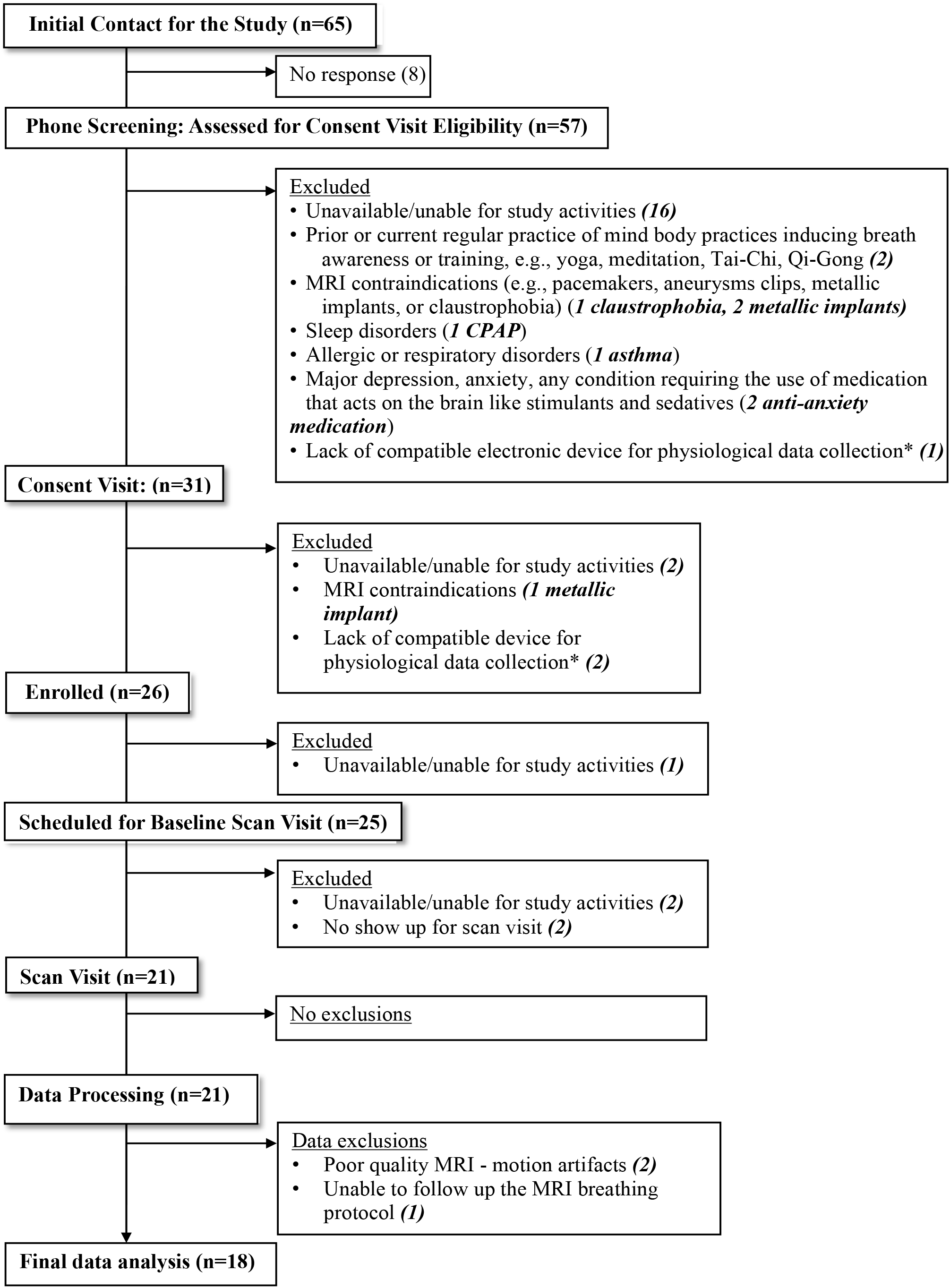
Study Flow Chart. *Our study utilized physiological data devices to objectively track participants’ home practice during the 8-week interventions. We excluded participants who did not have a compatible electronic device such as smartphone or tablet (see Table S1).

**Table 1.** Study group characteristics.

| *N* | Age range | Age in years<br>Mean (SD) | Sex | BMI<br>Mean (SD) | BP<br>Mean (SD) | Race | Ethnicity |
| --- | --- | --- | --- | --- | --- | --- | --- |
| 18 | 18-61 | 34.9 (14) | F, 8<br>M, 10 | 24.2 (5.6) | Systolic, 123 (17)<br>Diastolic, 75 (16) | 1 African American<br>4 Asian<br>1 More than one race<br>1 Unknown<br>11 White | 2 Hispanic or Latino<br>16 Not Hispanic or Latino |

### 2.2. Experimental Methodology

Each subject’s imaging visit lasted approximately 3-hours including study instructions, 1-hour MRI scans, and a set of questionnaires (as part of the RCT activities; not reported herein). Upon arrival for their imaging visit, we measured each subject’s temperature, blood pressure, and height and weight for body mass index (BMI). We then transitioned subjects to a mock scanner room for a ~30-min instruction for the breathing practices to be performed during the RT-PCMRI scans. We first explained and demonstrated each breathing practice, then guided subjects to perform at their own pace first seated on a chair, and then in supine in the mock scanner to mimic the MRI environment.

#### 2.2.1. MRI Breathing Protocol

We instructed subjects to perform the following breathing protocol first in the mock scanner for training purposes, and then in the MRI instrument during the ~1-minute RT-PCMRI measurements, each to be collected twice: (1) <u>spontaneous breathing</u> (SponB), (2) <u>slow breathing</u> (SlowB), (3) <u>deep abdominal breathing</u> (DAB), (4) <u>deep diaphragmatic breathing</u> (DDB) (5) <u>deep chest breathing</u> (DCB). See **Table S2** for the breathing protocol details.

#### 2.2.2. Rationale for the MRI Breathing Protocol Design

We chose breathing practices that were easily performed in supine in an MRI instrument without any constraints, and were less likely to cause head motion artifacts. We began with spontaneous breathing to observe each subject’s unique resting-state (natural) breathing patterns, and corresponding instantaneous CSF velocity waveforms. Since a key principle of a regular yogic breathing practice -that have been associated with health-benefits^57,58,62^ −is to make the breath slower and deeper, we included slow and deep breathing practices that were likely to have immediate impact on pulsatile CSF motion, and create larger changes compared to spontaneous breathing in magnitude and frequency of pulsatile CSF motion based on our pilot studies and literature review ^12–14^. This would allow us (i) to compare each subject’s unique spontaneous versus yogic breathing patterns, and corresponding CSF velocity waveforms (ii) to then quantify changes in magnitude and frequency components of CSF for identifying the primary driving force of CSF (respiratory versus cardiac components) during spontaneous versus yogic breathing practices.

#### 2.2.3. Breathing Rate and Depth

At the core of yogic breathing practices lies awareness and training of the breath. We designed our RCT yogic breathing interventions from Raja Yoga^63^ practices in the Himalayan Tradition, in which yogic breathing is suggested to be performed within each person’s own capacity for safety reasons, with inhale/exhale to be extended and expanded with caution through regular long-term practice. With that goal in mind, for the MRI breathing protocol, we specifically avoided enforcing any specific rate or depth for inhale/exhale other than giving a choice of rate (e.g., 3 to 5 counts with a count rate of 1/sec). This approach would allow us (i) to quantify changes in each subject’s unique pre-versus post-intervention breathing patterns, thus CSF waveform patterns, (ii) to then determine whether a regular long-term practice would influence the ability and capacity of subjects to modulate their breathing patterns (e.g., slower, deeper, rhythmical, smooth transitions between inhale/exhale), thus corresponding CSF dynamics.

#### 2.2.4. Subject preparation in the MR instrument

After being introduced to the breathing techniques in the mock scanner, we transitioned subjects to a 3T MRI instrument (MAGNETOM Prisma, Siemens Healthineers, Erlangen, Germany) for baseline data acquisitions using a 64-channel head and neck coil. We positioned subjects in supine, and provided them with (i) a bolster placed under the knees, (ii) foam pads under the elbows, (iii) pads around head and neck for comfort and minimizing motion artifacts, (iv) blankets for warmth, (vi) a wireless finger pulse sensor (Siemens Health) for pulse data collection, and (vii) a respiration bellow (Siemens Health) for respiration data collection during the entire RT-PCMRI data acquisitions. We instructed subjects to lie still in supine during the entire data acquisition.

### 2.3. Data Acquisition

We utilized a 1-hour data acquisition protocol, similar to our previous work^14^, consisting of anatomical MRI acquisitions, followed by simultaneous recordings of our previously established RT-PCMRI ^14^ acquisition, respiration and finger pulse acquisitions. Briefly, for consistency across all subjects, we aimed to measure CSF at an angle perpendicular to the spinal cord at the level of the foramen magnum (FM) (**Fig. 2 A_1-2_** green lines). To determine the location of FM, we first collected anatomical MR images using a T_2_-weighted fast spin echo (HASTE, repetition time (TR) 1200 ms, echo time (TE) 80 ms; **Fig. 2A_1_**); a 3D T_1_-weighted gradient echo sequence (MPRAGE; TR 2300 ms; TE 2.32ms; **Fig. 2A_2_**). We then acquired a cardiac gated PCMRI (TR 26.4 ms, TE 9.04 ms; **Fig. 2A_3_**) prior to RT-PCMRI to ensure proper slice location and angle for visibility of CSF pulsations, and that CSF was not obstructed. Upon confirming the pulsatile CSF motion (**Fig. 2A_3_**), we then acquired ~1-minute RT-PCMRI (**Fig. 2A_4-5_**) at the same slice location and angle when subjects performed each of the breathing practices. RT-PCMRI sequence parameters included: velocity encoding value (VENC) 5 cm/s, temporal resolution ~55ms, flip angle 30 degrees, matrix size 78 ×128, field of view (FOV) 196 × 323 mm (in-plane resolution ~ 2.5 × 2.5 mm), EPI factor 7, slice thickness 10 mm, TR 108.88 ms, TE 8.74 ms). RT-PCMRI has previously been described in detail ^14^. During the RT-PCMRI acquisitions, we simultaneously collected respiration and pulse data with a sampling frequency of f_s_=400 Hz.

**Figure 2.**
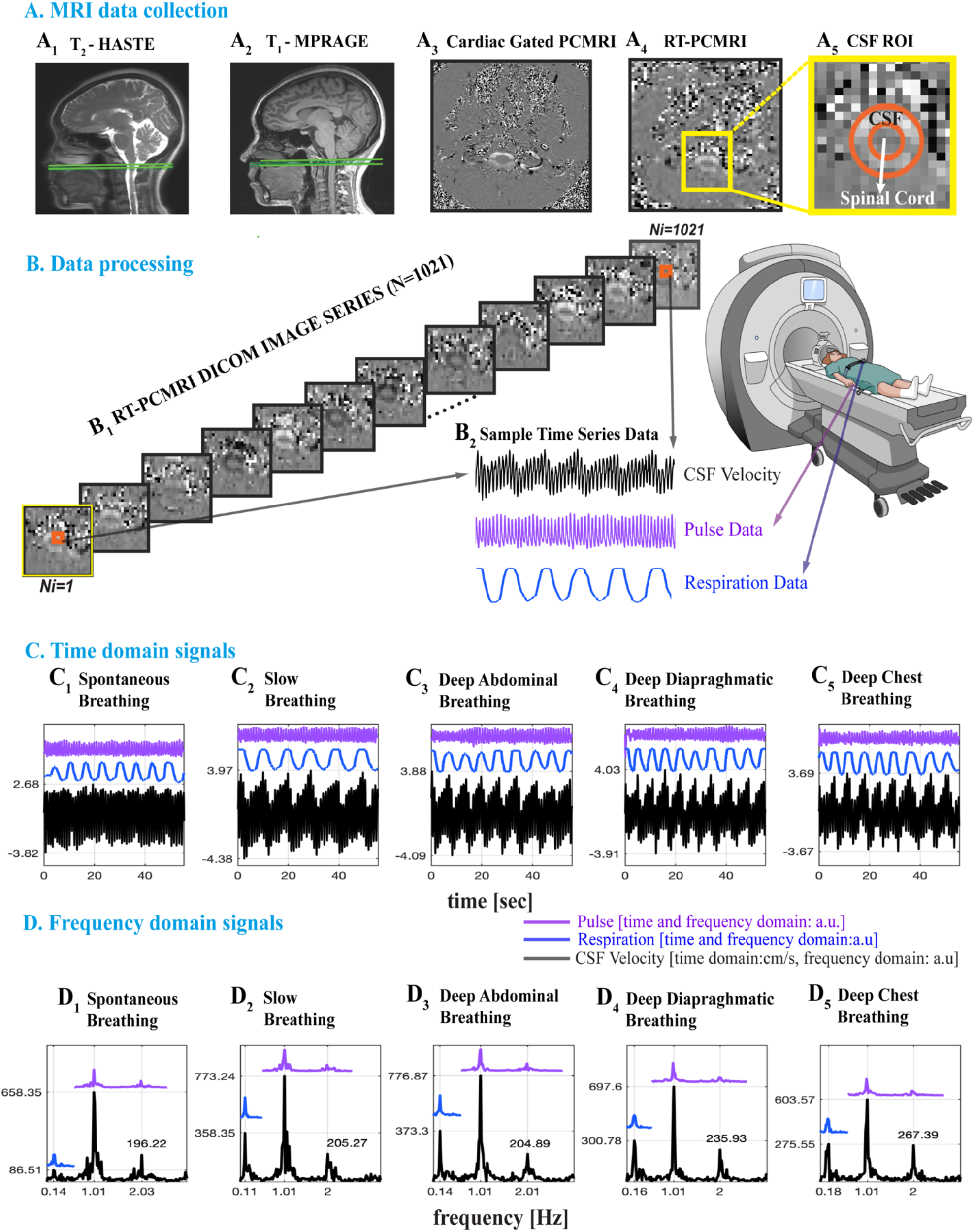
**A_1-2_** Sagittal anatomical MRI images showing the CSF measurement location at the level of foramen magnum (green lines) of a 37-year-old female. **A_3_** Axial images for cardiac gated PCMRI and RT-PCMRI velocity distribution. Cardiac-gated PCMRI is first collected for confirming CSF pulsation visibility prior to RT-PCMRI. ~1-minute RT-PCMRI is then collected at the same location during five breathing patterns. **A_5_** A detailed image of region delineates the voxels of CSF and spinal cord (orange lines) and surrounding tissue. **B_1_** RT-PCMRI DICOM phase images (N=1021) are collected for each breathing pattern repeated twice, resulting a total of 214,410 images processed for the n=21 subjects, and 183,780 images utilized for the results of n=18 subjects. **B_2_** Sample time series of CSF from single voxel RT-PCMRI (2.5 mm × 2.5 mm). Respiration and pulse data were simultaneously recorded, and temporally registered with the RT-PCMRI time series. **C-D** Time and frequency domain analysis of five breathing conditions: spontaneous breathing (SponB), slow breathing (SlowB), deep abdominal breathing (DAB), deep diaphragmatic breathing (DDB), and deep chest breathing (DCB) (with the last three forming a specific yogic breathing technique called *three-part breath*; see Table S2). When compared to SponB, both time domain maximum (positive; cranially directed) instantaneous CSF velocity values (in C), and peak respiration frequency amplitudes (in D) increase during SlowB, DAB, DDB, and DCB.

### 2.4. Data Processing

Each RT-PCMRI acquisition series produced 2042 images (1021 magnitude and 1021 phase, **Fig. 2B_1_**). In total (for N=21 subjects, five breathing conditions (SponB, SlowB, DAB, DDB, DCB) each repeated twice) we have acquired 428,820 RT-PCMRI (magnitude and phase) images, and processed the needed 214,410 RT-PCMRI phase images for obtaining CSF velocity time series. We have developed a semi-automated protocol for post-processing all MRI DICOM images, and respiration and pulse data time series using MATLAB software packages [2019-2020; Mathworks].

#### 2.4.1. CSF ROI and Velocity Waveforms

A common method ^30,32^ in conventional PCMRI studies to obtain CSF velocity time series is to average CSF across all voxels within the outlined region of interest (ROI), which may potentially cause spatial noise due to border zone partial volume effects^64^. Achieving a high temporal resolution for RT-PCMRI further may reduce the spatial resolution compared to conventional PCMRI, for which we previously developed a correlation mapping technique that allowed us to extract and average only highly correlated CSF voxels for an averaged CSF velocity time series. In this study, we are interested in obtaining and comparing true spatial and temporal CSF velocity values [cm/s] for each breathing practice. Therefore, to capture true spatial peak velocities to our best ability, we utilized a 2-step process to evaluate CSF velocity waveforms at a single voxel^9^ (**Fig. S1**): we first extracted highly correlated CSF voxels (greater than 0.7 correlation coefficient ^14^) with our previously developed correlation mapping technique ^14^, and then visually compared the CSF ROI voxels on RT-PCMRI images (spatial resolution of 2.5 × 2.5 mm) with the cardiac-gated PCMRI images (higher spatial resolution of 0.625 × 0.625 mm) to confirm the location of a single voxel of interest within CSF ROI. We are interested in maximum capacity of (participant) breathing impact on CSF. Anterior CSF velocities were usually larger than posterior velocities across our study population. We selected a single anterior voxel, within the CSF space, with greater velocity, which was usually among the highest correlated voxels (greater than 0.9 correlation coefficient) obtained from the correlation mapping technique. In addition, to confirm deep breathing practices did not cause artifacts in velocity values, potentially by B0 field changes in head caused by the motion of torso during deep breathing, we computed CSF velocities in a set of voxels within static tissue, and confirmed there were no respiratory or cardiac frequency components (**Fig. S2**).

#### 2.4.2. Time and Frequency Domain Signals of Interest

Previous studies ^9,13,14,37,47^ reported vasomotion, respiration, and cardiac (1^st^ harmonic) components of CSF signals. We observed (**Fig. S3**) higher order harmonics of cardiac pulsations in our preliminary analysis of frequency domain CSF velocity signals, which provides important information for determining the mechanisms, and their relative contribution to pulsatile CSF velocities. Having observed 1^st^ and 2^nd^ cardiac harmonics but not 3^rd^ or 4^th^ harmonics in all subjects, we have included 2^nd^ cardiac harmonics in our analysis. In short, we are interested in four distinctive CSF velocity time series: instantaneous (iCSF), respiratory (rCSF), and cardiac 1^st^ (c_1_CSF) and 2^nd^ harmonics (c_2_CSF). To remove higher frequency noise and observe only the rCSF, respiration, c_1_CSF and c_2_CSF, we low-pass filtered raw CSF velocity time series using a 4^th^ order Butterworth filter with a cut of frequency of 4 Hz, which provided time domain iCSF velocity signals (**Fig. 2C_1-5_**). We then computed frequency domain signals using a fast Fourier transform (**Fig. 2D_1-5_**).

To separate and investigate rCSF, c_1_CSF and c_2_CSF (**Fig. 3**), we first computed frequency bands of each of the three components for each subject and breathing condition. This allowed us to take into consideration of the individual and unique variations of frequency bands for accurate time domain velocity waveforms as well as frequency domain power calculations for each of the three components. We then filtered instantaneous CSF velocity waveforms (**Fig. 3A_1_**), using the individual frequency bands **(Fig. 3A_2_**), in (i) respiration frequency band of (estimated as typically f < ~0.6 Hz band), and (ii) cardiac 1^st^ harmonic frequency band (estimated as typically ~0.6 < f < ~1.6 Hz band), and (iii) in cardiac 2^nd^ harmonic frequency band (estimated as typically ~1.6 < f < ~2.7 Hz band) **(Fig. 3A_3-5_**). We repeated the above procedure for each subject’s each breathing condition, and obtained time and frequency domain signals for all four distinctive CSF velocity waveforms.

**Figure 3.**
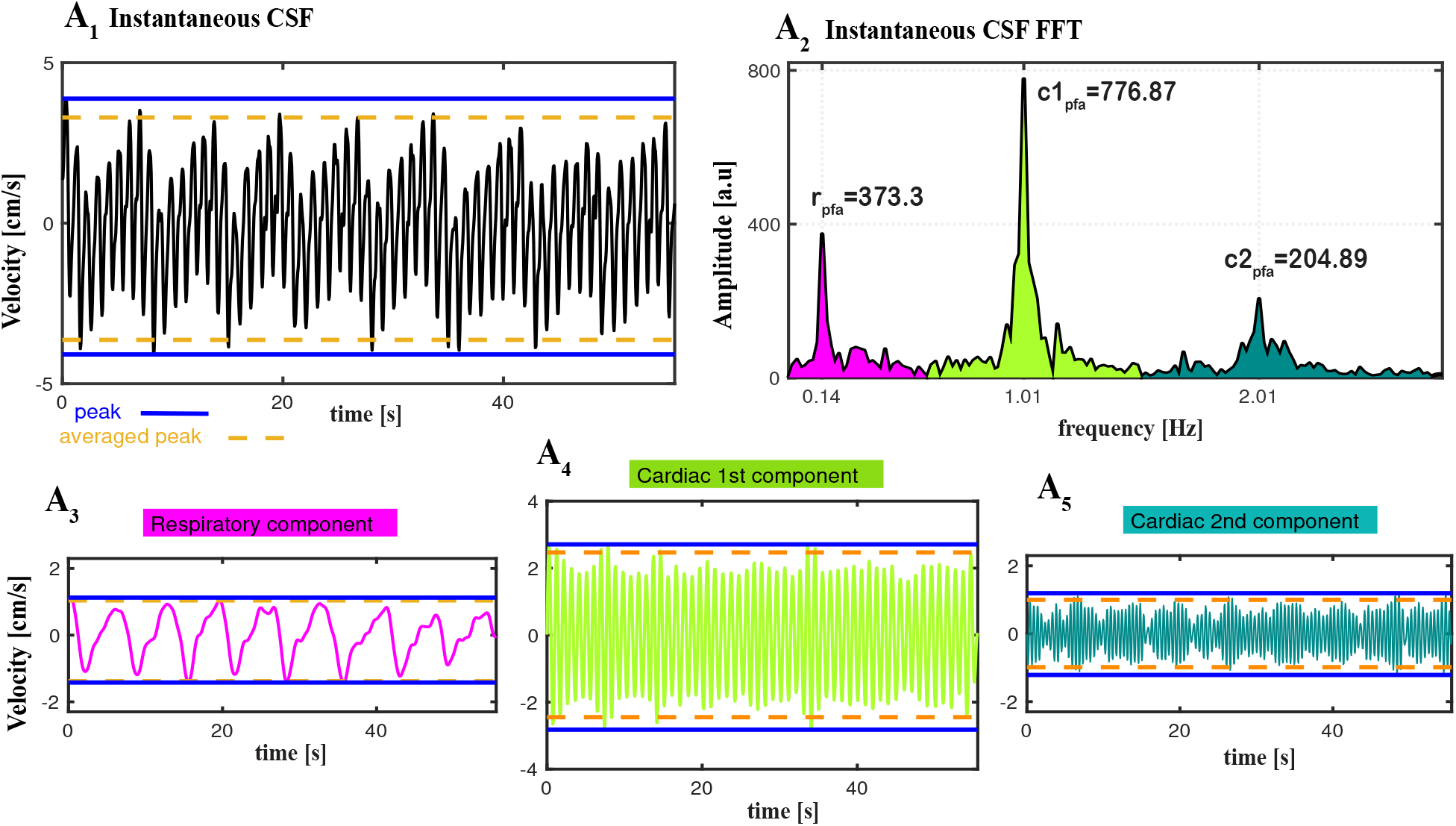
Separation of instantaneous CSF (iCSF) velocity waveforms (measured during deep abdominal breathing (DAB) for a 37 y-o female) into three components : respiratory (rCSF), cardiac 1^st^ harmonic (c_1_CSF) and cardiac 2^nd^ harmonic (c_2_CSF). **A_1_** Time domain iCSF velocity waveforms. **A_2_** Frequency domain iCSF obtained from Fast Fourier Transform (FFT) of iCSF velocity time series presenting peak frequency amplitudes for respiration (r_pfa_), cardiac 1^st^ harmonic (c_1pfa_) and 2^nd^ harmonic (c_2pfa_). We filtered iCSF within the bandwidth of each component - respiratory (magenta), cardiac 1^st^ harmonic (light green) and 2^nd^ harmonic (dark green) - in order to obtain and compare the characteristics of individual pulsatile velocity time series **A_3_** rCSF, **A_4_** c_1_CSF, and **A_5_** c_2_CSF. We computed maximum (positive; cranially directed) and minimum (negative; caudally directed) CSF velocity value in two ways: (i) peak value obtained as the highest and lowest point (blue solid line) during the entire time series, and (ii) averaged peak value (orange dash line) obtained by averaging the local maximums (and minimum, respectively) above a threshold set at 97.5^th^ percentile (and 2.5^th^ percentile, respectively). We used averaged peak values in statistical analysis to reduce temporal noise due to any transient events that may cause abrupt peaks (e.g., unexpected deep sigh) or lower peaks (indicating less than maximum capacity) caused by “subject fatigue” during deep breathing conditions.

In parallel, to confirm that estimated frequencies of CSF signals match with the physiological data, we filtered respiratory sensor and pulse sensor data in the same frequency band of respiratory and cardiac components of CSF velocity waveforms. For visualization purposes, we arbitrarily scaled the respiration and pulse data to compare with CSF velocity waveforms (**Fig. 2C-D_1-5_**). We confirmed the respiratory component in respiration data, and cardiac (1^st^ and 2^nd^) harmonic components in pulse data (blue and purple lines in **Fig. 2D_1-5_**).

### 2.5. CSF Metrics

#### 2.5.1. Time Domain CSF Metrics

We computed the following metrics from time domain CSF velocity waveforms iCSF, rCSF, c_1_CSF, c_2_CSF for each subject and breathing condition: 1) peak^13,14,37^ maximum (cranially directed value at the highest point) and peak minimum (caudally directed at the lowest point) during ~1-minute time series; 2) averaged peak maximum and averaged peak minimum obtained from the average of local peak maximums (and peak minimums, resp.) above (and below, resp.) a threshold set at 97.5^th^ percentile ^65,66^ (and 2.5^th^ percentile, resp.); 3) range^32^ of peak maximum to peak minimum; 4) range of averaged peak maximum to averaged peak minimum; 5) displacement computed from the integration of the CSF velocity time series [cm/s] and converted to [mm]. Lastly, we computed % change in these metrics, from SponB to SlowB, DAB, DDB, and DCB.

Note that the traditional method for computing cranially- and caudally-directed velocities are to compute peak maximum and peak minimum values. In addition to peak maximum and minimum, for this study, we also computed averaged peak maximum and averaged minimum values. Since our goal during each breathing condition is to capture true maximum capacity of CSF velocity, the use of averaged peak approach allowed us to reduce temporal noise caused by i) random transient events that are not part of the regular breathing pattern (e.g., unexpected deep sigh) resulting in greater peak values, or ii) “participant fatigue” experienced during performing slow and/or deep breathing conditions resulting in lower peak values. See **Fig. 3** blue and orange dash lines for a comparison of peak versus averaged peak values.

#### 2.5.2. Frequency Domain CSF Metrics

From the instantaneous CSF velocity waveforms, we have computed 1) peak frequencies, and 2) peak frequency amplitudes (**Fig. 3A_2_**). Additionally, to observe individual peak frequency amplitude ratio changes, we computed 3) a peak-to-peak frequency amplitude ratio [r/c_1peak_] and [r/c_2peak_] calculated as the ratio of rCSF peak frequency amplitude to the c_1_CSF, and c_2_CSF peak frequency amplitudes in the frequency domain (**Fig 4.A_1-5_**).

**Figure 4.**
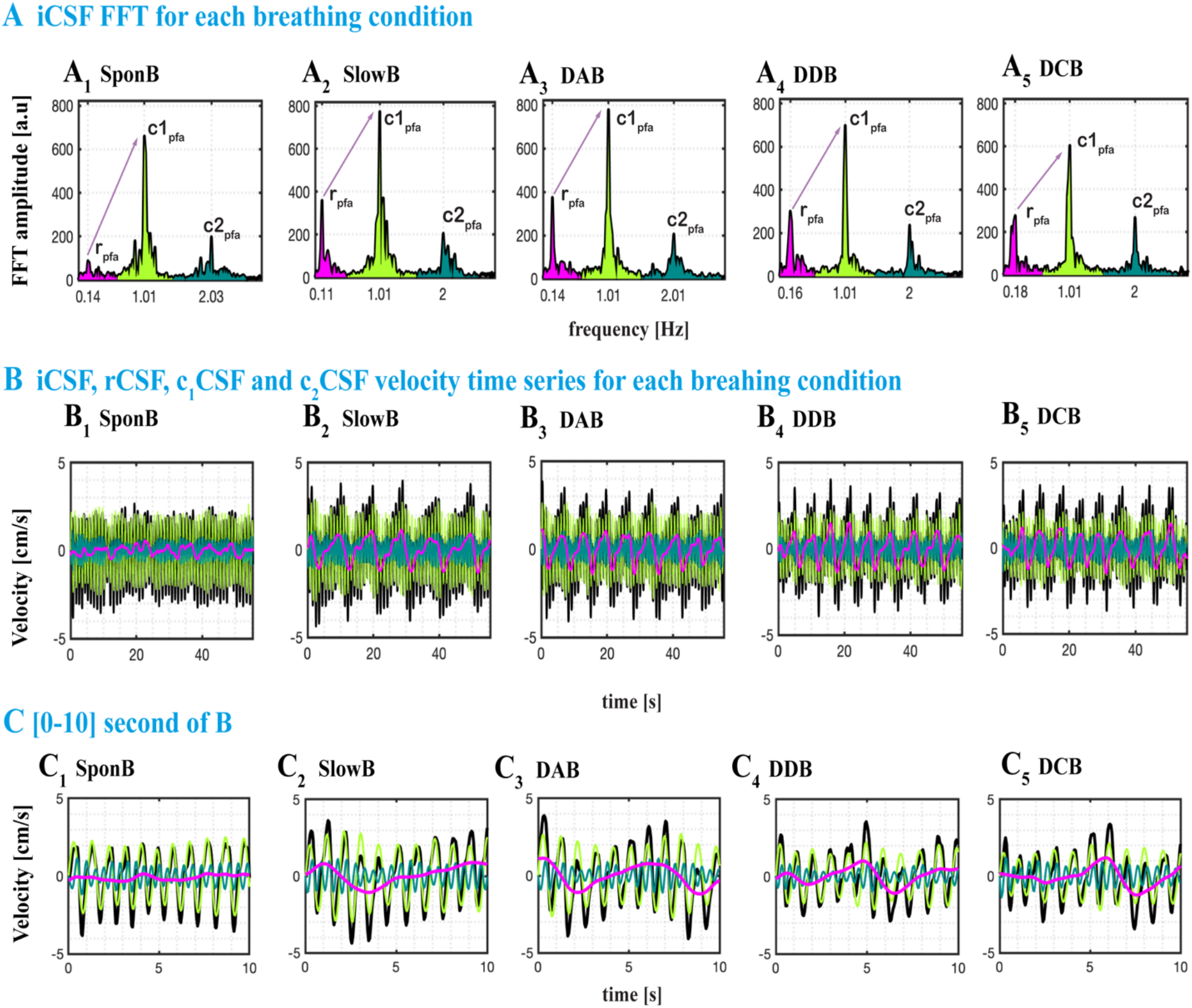
Sample datasets from a 37 y-o female presenting all four distinctive CSF signals (black: iCSF, magenta: rCSF, light green: c_1_CSF, dark green: c_2_CSF) during SponB, SlowB, DAB, DDB, and DCB. **A_1-5_** Frequency domain iCSF signals presenting the changes in peak frequencies [x-axis: r_pf_, c_1pf_, and c_2pf_; Hz] and peak frequency amplitudes [y-axis; r_pfa_, c_1pfa_, and c_2pfa_; a.u]. There was a decrease in respiration peak frequency during SlowB compared to SponB, and increase in respiration peak frequency amplitude r_pfa_ during all four yogic breathing techniques (A_2-5_) due to increased respiratory movement. We computed peak frequency amplitude ratios [r/c_1peak_] and [r/c_2peak_] (e.g., [r/c_1peak_] indicated by sample purple arrows) for observing the changes, and power ratios [r/c_1power_] and [r/c_2power_] for testing the relative contribution of each component to instantaneous CSF. **B_1-5_** Time domain CSF signals presenting an increase in both cranially directed iCSF and rCSF velocities during four yogic breathing techniques compared to SponB; with detailed waveforms presented in [0-10] s time window in **C_1-5_**.

#### 2.5.3. Relative contribution of rCSF, c_1_CSF, c_2_CSF Signals

To compare the contribution of rCSF, c_1_CSF, and c_2_CSF and determine the primary regulatory force(s) for pulsatile CSF, we computed 1) estimated frequency band of rCSF, c_1_CSF, and c_2_CSF (**Fig. 3A_2_**, **Fig 4.A_1-5_**), 2) power of rCSF, c_1_CSF, and c_2_CSF (defined^14^ as the integral of the square of the amplitude spectrum over the corresponding frequency band), and 3) relative contribution^13,14,37^ of the respiration versus cardiac components by defining power ratio^14^ [r/c_1power_] and [r/c_2power_] calculated as the ratio of the power of the rCSF to the power of c_1_CSF and c_2_CSF, respectively.

### 2.6. Statistical Analysis

To test the differences between spontaneous and four yogic breathing techniques, we used mean and standard deviation (SD) for the following time and frequency domain CSF metrics (i) averaged peak maximum and minimum values for iCSF, rCSF, c_1_CSF, c_2_CSF; (ii) range of iCSF averaged peak maximum and minimum values, (iii) iCSF displacement, (iv) peak frequencies, (v) peak frequency amplitudes, (vi) power values for rCSF, c_1_CSF, c_2_CSF, (vii) frequency peak-to-peak amplitude ratios, and (viii) power ratios for a total of 23 metrics (**Table 2**).

**Table 2.**
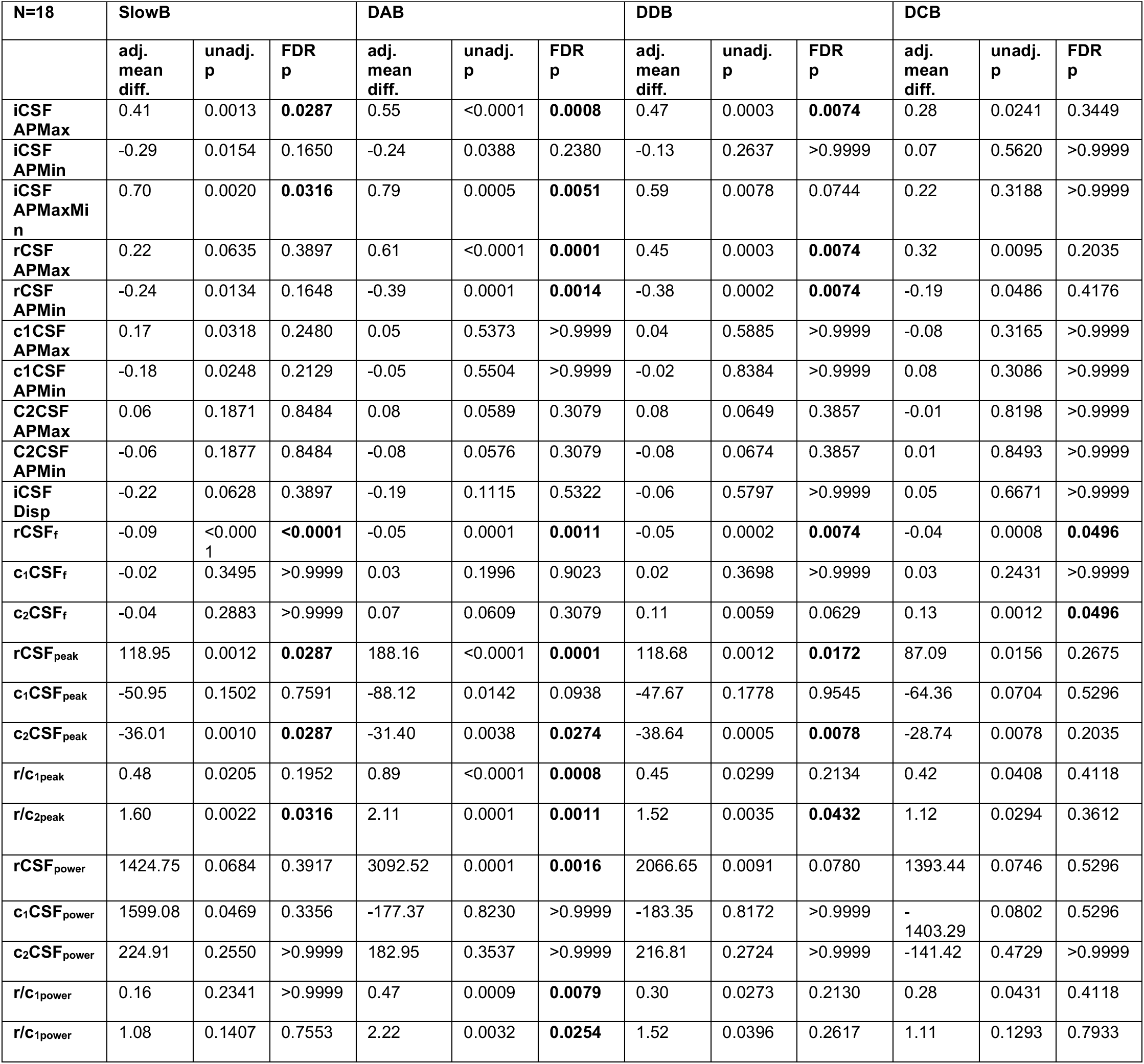
Results of statistical testing, implemented for differences between SponB and four yogic breathing techniques. SponB: spontaneous breathing, SlowB: slow breathing, DAB: deep abdominal breathing, DDB: deep diaphragmatic breathing, DCB: deep chest breathing, iCSF: instantaneous-CSF, rCSF: respiratory-CSF, c_1_CSF: cardiac 1^st^ harmonic CSF, c_2_CSF: cardiac 2^nd^ harmonic CSF, APMax: averaged peak maximum, APMin: averaged peak minimum, APMaxMin: averaged peak maximum to averaged peak minimum, Disp: displacement, rCSFf: peak frequency of rCSF, c_1_CSFf: peak frequency of c_1_CSF, c_2_CSFf: peak frequency of c_2_CSF, rCSFpeak: peak frequency amplitude of rCSF, c_1_CSFpeak: peak frequency amplitude of c_1_CSF, c_2_CSFpeak: peak frequency amplitude of c_2_CSF, r/c_1peak_: peak frequency amplitude ratio of rCSF to c_1_CSF, r/c_2peak_: peak frequency amplitude ratio of rCSF to c_2CSF_, rCSFpower: power of rCSF, c_1_CSF_power_: power of c_1CSF_, c_2_CSF_power_: power of c_2CSF_, r/c_1power_: power ratio of rCSF to c_1_CSF, r/c_2power_: power ratio of rCSF to c_2_CSF.

Independent statistical analysis was conducted by A.H. in R version 4.0.3 (R Core Team, Vienna, Austria). Data were visually inspected, and extreme values were double checked and remained in the data if confirmed by the principal investigator to be reasonable values. Mixed effects linear regression models were built to analyze the associations between each outcome measure and each of the four experimental breathing conditions (SlowB, DAB, DDB, and DCB). Each model included a random subject effect to characterize within-person correlations over repeated measures. Normality assumptions of each model were checked by visual inspection of Q-Q Plots. For multiple comparisons, type I error rate was controlled by using the Benjamini-Yekutieli false detection rate (FDR) procedure ^67^, with an overall FDR of 0.05, using the R program “p.adjust”. Unadjusted p-values and FDR corrected p-values are provided in **Table 2,** and p-values mentioned within the text are FDR corrected p-values.

In addition, associations between demographic covariates and outcomes were inspected visually, and tested for significance by Spearman’s rank-order correlation (continuous covariates) or t-test (dichotomous covariates) if a possible association was seen. We reported associations found to be significant in section 3.5. However, the regression models of the main analyses were not adjusted for these covariates in order to maintain consistency between models and so as not to overfit the models.

## 3. Results

### 3.1. Baseline Group Characteristics

Twenty-one subjects completed the baseline study activities. We processed 214,410 RT-PCMRI phase images, and removed datasets from final data analyses for three subjects; two subjects due to motion artifacts (e.g., compromised image quality), and one subject due to inability to follow the MRI breathing protocol. We then utilized 183,780 RT-PCMRI phase images from 18 participants for the final data analyses. See **Fig. 1** for study flow chart, and **Table 1** for baseline group characteristics.

We presented sample datasets from a set of participants to demonstrate the changes in time domain and frequency domain CSF metrics during SponB versus yogic breathing (**Fig. 2**, and **Fig. 4–5)**. Group summary metrics (N=18; all five breathing conditions) used for statistical analysis (obtained in 2.6) are shown in **Table S3-4**, providing [mean, SD, %Δ] and statistical analysis results in **Table 2** providing [adjusted mean difference, unadjusted p-value, FDR p-values]. Since we had 23 outcome measures to test the differences, we chose FDR which utilizes a Benjamini-Yekutieli algorithm, equipped to handle dependence between multiple outcome measures. We presented all group summary CSF metrics (obtained in 2.5.1-3) in **Table S3-5**, and main findings for SponB versus DAB in **Fig.6**. We will discuss group summary statistics in the following sections.

**Figure 5.**
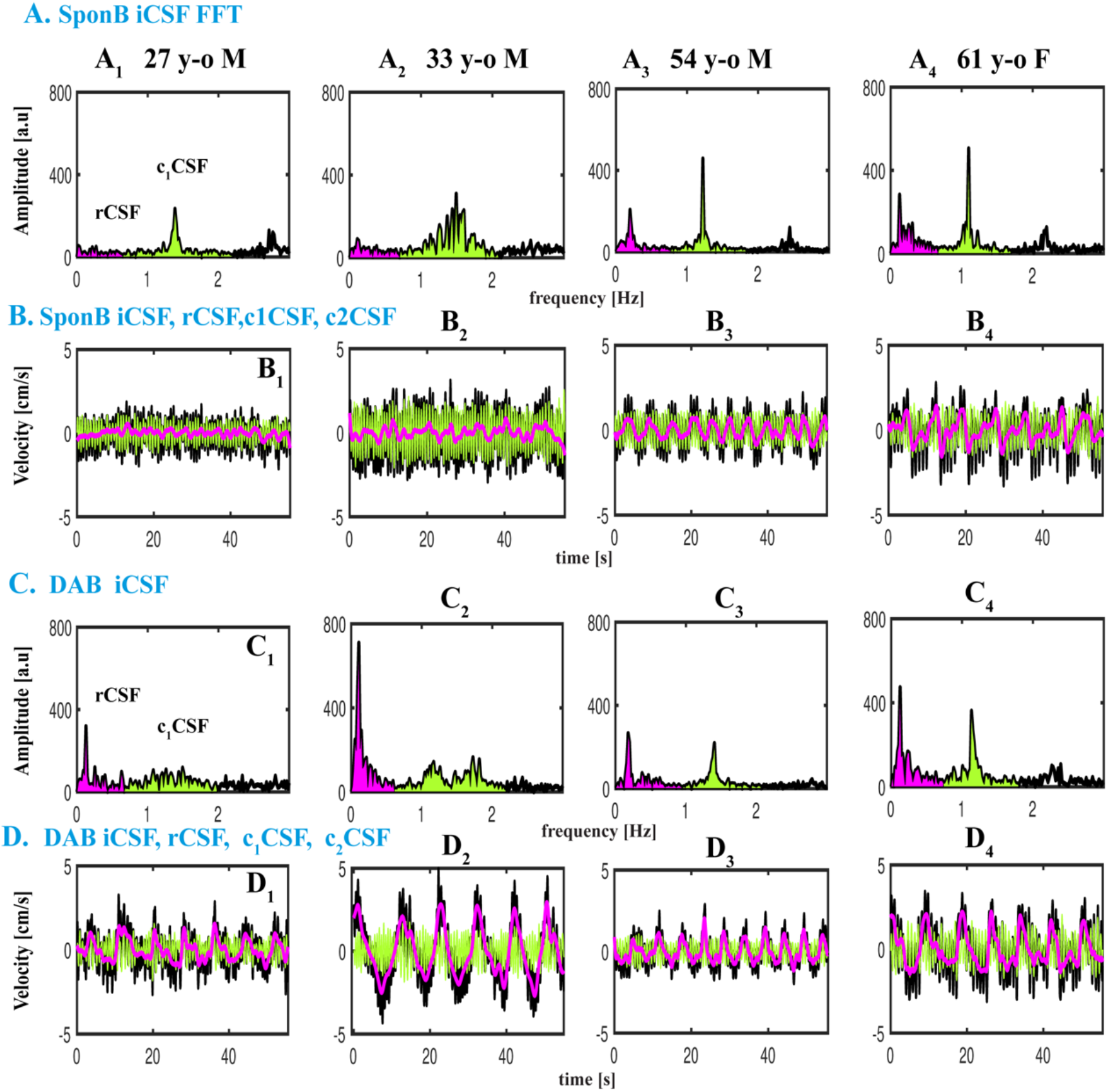
Datasets from four participants during SponB versus DAB to demonstrate the relative contribution of respiration versus cardiac 1^st^ harmonic to pulsatile CSF, which was computed using a power analysis. **A** SponB (and **C** DAB) frequency domain iCSF signals. **B** (and **D**) Time domain velocity time series for iCSF (black), rCSF (magenta), c_1_CSF (lightgreen), and c_2_CSF (dark green). During SponB, across the 18 participants, cardiac pulsation was the major driver for pulsatile CSF, including four participants presented in A-B. During DAB, while across the 18 participants, there was a comparable contribution of cardiac 1^st^ harmonic and respiration, for these four participants, respiration was the major driver for pulsatile CSF due to significantly increased breathing depth resulting in increased [r/c_1power_] (A vs. C). Also see increase in cranially directed iCSF and rCSF peak velocities (B vs. D).

**Figure 6.**
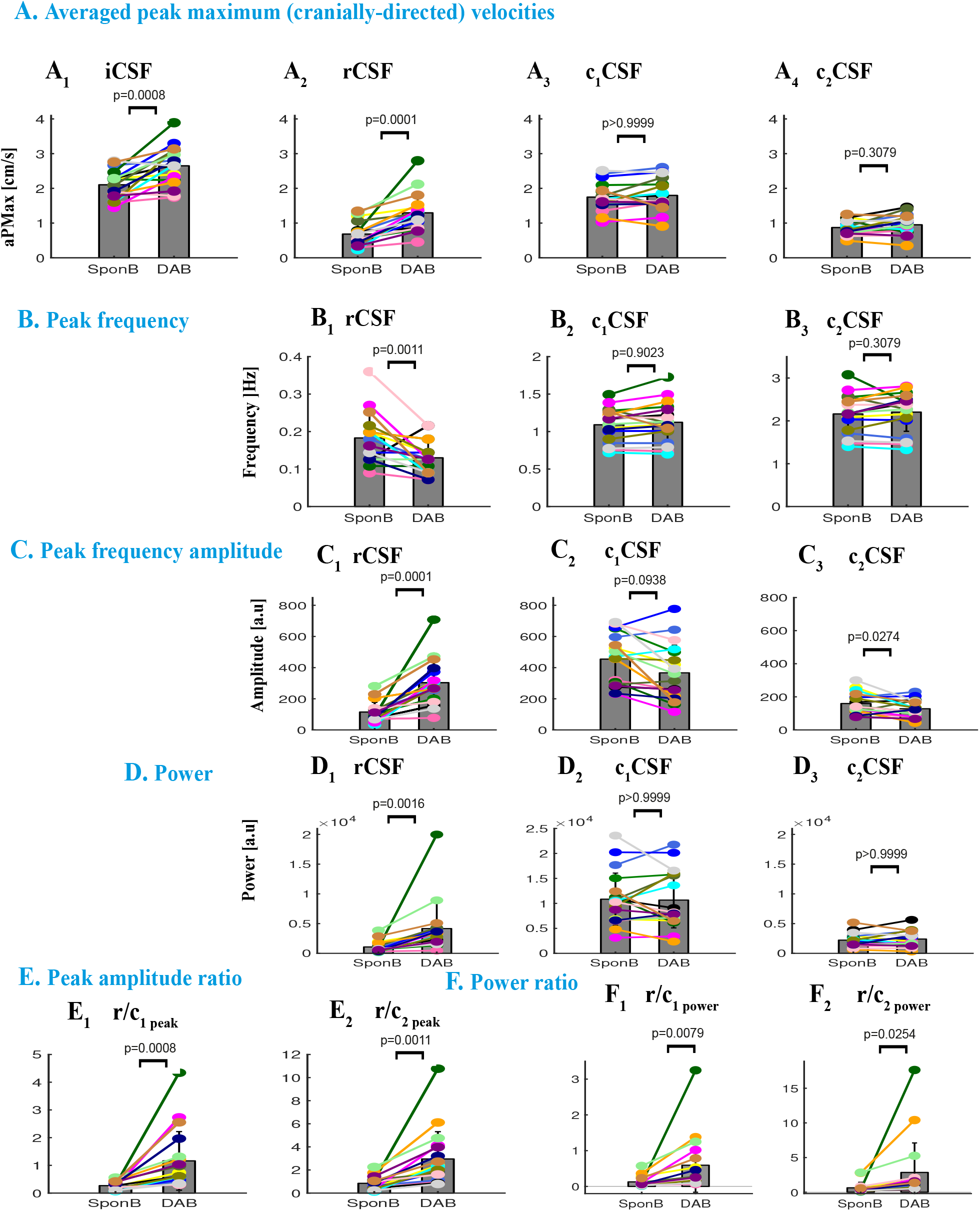
Comparison of time and frequency domain metrics used for SponB versus DAB. **A** Time domain: CSF velocity averaged peak (maximum cranially-directed) values for iCSF and rCSF significantly increased (p=0.0008, and p=0.0001), but not for c1CSF and c2CSF. **B-F** Frequency domain: **B** Peak frequency for rCSF significantly decrease (p=0.0011), with no significant changes for c_1_CSF and c_2_CSF. **C** Peak frequency amplitudes significantly increased for rCSF (p=0.0001), and significantly decreased (p=0.0274) for c_2_CSF with no significant changes for c1CSF. **D** Power for rCSF significantly increased (p=0.0016) with no significant changes for c_1_CSF and c_2_CSF. **E** Peak amplitude ratios [r/c_1peak_] and [r/c_2peak_] significantly increased (p=0.0008, and p=0.0011). **F** Power ratios [r/c_1power_] and [r/c_2power_] significantly increased (0.0079, and p=0.0254), with a greater contribution of respiration compared to cardiac 1^st^ harmonic for four participants (as shown in Fig. 5).

### 3.2. Changes in Time Domain CSF Metrics During SponB versus Yogic Breathing

For all subjects (N=18; **Table 2** and **Table S3**), during five breathing conditions, cranially-directed velocities (averaged peak maximum) were iCSF [2.10 to 2.65] cm/s, rCSF [0.68 to 1.29] cm/s, c_1_CSF [1.67 to 1.92] cm/s, and c_2_CSF [0.86 to 0.95] cm/s resulting in greater c_1_CSF velocities and comparable velocities for rCSF and c_2_CSF. (See **Table S3** for mean and SD values during each breathing condition). When comparing the cranial iCSF velocities for SponB versus yogic breathing, we found an increase of 16% - 28% in cranial iCSF velocities during all yogic breathing conditions with statistical significance for SlowB (22%, p=0.0287), DAB (28%, p=0.0008; **Fig. 6A_1_**), DDB (23%, p=0.0074), and an increase of 60% - 118% in cranial rCSF velocities during all yogic breathing conditions with statistical significance for DAB (118%, p=0.0001, **Fig. 6A_2_**) and DDB (84%, p= 0.0074).

Caudally-directed velocities (averaged peak maximum) were iCSF [−2.67 to −3.03] cm/s, rCSF [−0.67 to −1.07] cm/s, c_1_CSF [−1.69 to −1.95] cm/s, and c_2_CSF [−0.87 to −0.95] cm/s resulting in greater c_1_CSF velocities and comparable velocities for rCSF and c_2_CSF. When comparing the caudal directed CSF velocities, we found an increase of 2% - 11% in caudal iCSF which did not reach statistical significance, and a decrease of 43% - 78% in caudal rCSF velocity with statistical significance for DAB (78%, p=0.0014) and DDB (68%, p= 0.0074). There were no statistically significant findings for cranial (**Fig. 6A_3-4_**) and caudal c_1_CSF and c_2_CSF velocities, as well as iCSF displacement during SponB versus yogic breathing.

### 3.3. Changes in Frequency Domain CSF Metrics During SponB versus Yogic Breathing

When compared to SponB (**Table 2** and **Table S4**), we found (1) a statistically significant decrease 18% - 42%; (p<0.05) in estimated rCSF peak frequency (respiration rate) during all yogic breathing conditions with most significance for SlowB (42%, p<0.0001) (see **Fig. 6B_1_** for DAB), (2) an increase of 101% - 234% in rCSF peak frequency amplitude with statistical significance for SlowB (141%, p=0.0287), DAB (234%, p=0.0001, **Fig. 6C_1_**), and DDB (160%, p =0.0172), (3) a decrease of 13% - 21% in c_2_CSF peak frequency amplitude with statistical significance for SlowB (20%, p=0.0287), DAB (15%, p=0.0274, **Fig. 6C_3_**), and DDB (21%, p=0.0078). There were no statistically significant changes in peak frequency for c_1_CSF and c_2_CSF (**Fig. 6B_2-3_**), except an increase for DCB; 6%, p=0.0496, and peak frequency amplitude for c_1_CSF (**Fig. 6C_2_**).

Additionally, we found an increase of 158% - 359% in peak amplitude ratio or rCSF to c_1_CSF [r/c_1peak_] with statistical significance for DAB (359%, p=0.0008, **Fig. 6E_1_**), and an increase of 166% - 350% in [r/c_2peak_] with statistical significance for SlowB (223%, p=0.0316), DAB (350%, p=0.0011, **Fig. 6E_2_**), and DCB (265%, p=0.0432).

### 3.4. Relative contribution of rCSF, c_1_CSF, c_2_CSF During SponB versus Yogic Breathing

During yogic breathing compared to SponB (**Table 2** and **Table S4**), we found an increase of 187% - 472% in rCSF power with statistical significance for DAB (472%, p=0.0016, **Fig. 6D_1_**), and no statistically significant findings for c_1_CSF and c_2_CSF power (**Fig. 6D_2-3_**). We computed relative contribution of rCSF versus c_1_CSF and c_2_CSF using the power ratios [r/c_1power_] and [r/c_2power_]. Power ratio [r/c_1power_] for each breathing condition was [SponB; 0.13 ± 0.15], [SlowB; 0.29 ± 0.51], [DAB; 0.59 ± 0.78], [DDB; 0.43 ± 0.56] and [DCB; 0.40 ± 0.60] demonstrating cardiac 1^st^ as major source of pulsatile CSF during SponB. There was an increase of 248% - 534% in [r/c_1power_] during yogic breathing compared to SponB, with statistical significance for DAB (534%, p=0.0079, **Fig. 6F_1_**) when there was a comparable contribution of respiration and cardiac 1^st^ harmonic to pulsatile CSF. For instance, four of the 18 participants (**Fig. 5**) presented greater respiratory power compared to cardiac 1^st^ harmonic power during DAB versus SponB resulting in respiration as the major driver for pulsatile CSF during DAB for these four participants.

Power ratio [r/c_2power_] for each breathing condition was [SponB; 0.63 ± 0.81], [SlowB; 1.72 ± 3.52], [DAB; 2.85 ± 4.38], [DDB; 2.16 ± 2.96] and [DCB; 1.75 ± 2.39] demonstrating comparable contribution of respiration and cardiac 2^nd^ harmonic during SponB. There was an increase of 234% - 589% in [r/c_2power_] during yogic breathing compared to SponB, with statistical significance for DAB (589%, p=0.0254, **Fig. 6F_2_**).

### 3.5. Covariates of age, sex, BMI

We tested associations between demographic covariates and outcomes during SponB versus yogic breathing. Under the DCB condition, there was a positive correlation between age and change scores (defined as change from the SponB condition) for cranial rCSF velocity (rho=0.69, p<0.001), rCSF frequency peak amplitude (rho=0.75, p<0.001), rCSF power (rho=0.79, p<0.001), and [r/c_1peak_; rho=0.69, p=0.001], [r/c_2peak_; rho=0.67, p=0.002], [r/c_1power_; rho=0.80, p<0.001]. [r/c_2power_; rho=0.80, p<0.001]. There was also an association between sex and c_2_CSF frequency peak amplitude for the change score between SponB to DAB (p=0.009), and SponB to DCB (p=0.048), with a mean increase in peak amplitude for females and a mean decrease for males.

In short, when compared to SponB, the main results were as follows; there was (1) a statistically significant decrease in respiration rate 18% - 42% during yogic breathing with most significance for SlowB (42%, p<0.0001), (2) increase of 16% - 28% in cranially directed iCSF velocities with most statistical significance for DAB (28%, p=0.0008), with no significance for DCB, (3) in parallel, an increase of 101% - 234% in rCSF peak frequency amplitude with most significance for DAB (234%, p=0.0001), with no significance for DCB, (4) increase of 187% - 472% in rCSF power with statistical significance only for DAB (472%, p=0.0016), (5) increase of 248% - 534% in [r/c_1power_] and 234% - 589% in r/c_2power_ with statistical significance only for DAB (534%, p=0.0079, and 589%, p=0.0254, resp.), (6) positive association between age and change scores from SponB to DCB.

## 4. Discussion

To our knowledge, this study is the first to investigate the effects of a mind-body approach such as yogic breathing on CSF dynamics. We measured CSF velocities at the level of FM with a non-invasive RT-PCMRI approach, and found an immediate impact of four different types of yogic breathing techniques on pulsatile CSF velocities compared to spontaneous breathing. Results indicate the following findings (i) respiration rate significantly decreased during slow and deep yogic breathing techniques; (ii) cranial iCSF velocities and in parallel rCSF peak frequency amplitudes increased during yogic breathing with most statistical significance for DAB, and with no significance for DCB, (iii) cardiac pulsation was the primary driving force for pulsatile CSF during spontaneous breathing when there was a comparable contribution of rCSF versus c_2_CSF, and (iv) cardiac pulsation was the primary driving force for CSF during yogic breathing except during DAB when there is a comparable contribution of rCSF and c_1_CSF.

### 4.1. Mechanics of Respiratory CSF Dynamics

Using a respiratory bellow, we collected respiration data simultaneously with the RT-PCMRI, thus confirmed cranially-directed CSF during inhalation and caudally-directed CSF during exhalation. This result is in agreement with previous studies^24,39,68^ measuring CSF pressure recordings in response to respiratory changes, coughing and Valsalva maneuver, and non-invasive MRI studies^13,36,45^ investigating respiratory CSF velocities or flow volumes. Briefly, the transmission of venous pressure changes to the collapsible dura through thoracic and epidural veins lining the spine and around the vertebral column causes CSF movement in an ebb-and-flow manner. Lloyd et al^16^ recently showed respiratory CSF flow is driven by lumbar and thoracic spinal pressures, and that reduced intrathoracic pressure during inspiration draws venous blood from the lumbar spine and cranium towards the thorax.

Of the four yogic breathing conditions we used in our study (SlowB, DAB, DDB, and DCB), the three of them (SlowB, DAB, and DDB) significantly increased iCSF velocities, with most pronounced effects observed during DAB with no significant change during DCB. The difference between abdominal and chest (thoracic) breathing we observed is aligned with previous reports indicating abdominal breathing is associated with larger respiratory pressure changes compared to thoracic breathing^15,69^. Aktas et al ^15^ for instance recently demonstrated forced abdominal breathing-compared to forced thoracic breathing- has more pronounced effects on CSF movement within spinal subarachnoid space, resulting in upward net flow during both breathing patterns, whereas there were low flow rates in the cerebral aqueduct in both breathing patterns. They concluded that abdominal breathing was associated with larger CSF flow due to a more pronounced contraction of the diaphragm compared to thoracic breathing. Furthermore, they suggested that changes in CSF dynamics were due to changes in intrathoracic and intraabdominal pressure being transmitted to the epidural space through the paravertebral venous plexus.

There were no statistically significant changes between SponB and DCB in our study, which may suggest our study population primarily consisted of natural chest breathers although other explanations are possible. While we observed significant increase in cranial directed iCSF velocities during SlowB, DAB, DDB, there were no changes in caudal iCSF velocities suggesting exhalation during spontaneous and yogic breathing in our study population was passive^16^.

### 4.2. Primary sources of pulsatile CSF dynamics

The sources of pulsatile CSF velocity waveforms are cardiac pulsation, respiration and low frequency components such as vasomotion. Several studies recently examined primary regulator(s) of CSF movement and/or flow. For instance, (i) Dreha-Kulaczewski *et al* ^12^ presented CSF signal intensities (in arbitrary units) during forced inspiration, and suggested forced inspiration is the major driver of CSF while (ii) Takizawa et al.^37^ demonstrated velocities of cardiac-driven CSF at cerebral aqueduct were greater than respiratory-driven CSF, while displacement of respiratory-driven CSF was greater than cardiac-driven CSF, (iii) Mestre et al.^7^ more recently demonstrated cardiac pulsation is the primary regulator of CSF flow through perivascular spaces (PVSs) and is reduced in hypertension, and (iv) Fultz et al demonstrated CSF flow is driven by vasomotion during sleep.

In our study, we presented respiratory and cardiac components of CSF while separating the cardiac 1^st^ and 2^nd^ harmonic components. During spontaneous breathing, we found that the cardiac 1^st^ harmonic contributed greater power to pulsatile CSF velocities, with comparable contributions by respiration and the cardiac 2^nd^ component. This suggests that the cardiac 2^nd^ harmonic effect on pulsatile CSF dynamics is comparable to the effect of respiration.

Similarly, during yogic breathing, the cardiac 1^st^ harmonic contributed greater power to pulsatile CSF velocities, except during DAB, when there was a comparable contribution of respiration and cardiac 1^st^ harmonic effect. We found a decrease in c_2_CSF peak frequency amplitude for breathing conditions with significant increases in rCSF peak frequency amplitudes. During in all breathing conditions, we found a larger frequency amplitude of cardiac 1^st^ versus 2^nd^ harmonic, in agreement with earlier studies^70–72^ investigating intracranial pressure (ICP) measures and a recent study^73^ investigating CSF dynamics of the American alligator. To our knowledge, this is the first comprehensive report studying a higher order cardiac harmonic component in human CSF non-invasively, and during voluntarily controlled breathing conditions. Higher harmonics of CSF have not been well-documented. Wagshul et al. ^72^ for instance investigated higher cardiac-induced harmonics in ICP, and interpreted changes in brain pulsatility in the context of system compliance (of brain tissue, arterial, venous, and spinal thecal sac communication with brain through CSF spaces). Young et al.^73^ (i) studied variations of pulsatile CSF in Alligator in spinal canal and cranial cavity, (ii) found cardiac-induced harmonics in CSF (not above 3^rd^ order), (iii) hypothesized the absence of higher harmonics could be related to the reptilian meninges and compliance. Taken together, higher harmonics of CSF provide important information for determining the mechanisms regulating CSF dynamics, and need to be investigated in further studies.

Despite the significant increase in cranially directed iCSF velocities and in parallel in rCSF peak frequency amplitudes during SlowB, DAB, and DDB, cardiac pulsation was still the primary contributor (except during DAB), suggesting that the significant increase in CSF peak velocities or in rCSF peak frequency amplitudes did not necessarily mean that respiration was the major regulator for CSF. Thus, in future studies, we recommend doing a frequency domain power analysis to determine primary regulator(s) of pulsatile CSF dynamics. For instance, group summary results indicate that yogic breathwork increased both cranial directed CSF velocities and respiratory CSF peak amplitudes. However, only four individual subjects (**Fig. 5**) had greater respiratory power compared to cardiac power during DAB, suggesting (i) power contribution is critical, and (ii) respiration can be a major driver for pulsatile CSF dynamics depending on individual differences in breath “depth and location”. In short, even if CSF velocities may significantly increase with increased respiratory movement, if the increase in amplitude does not meet a certain threshold (e.g., not breathing deeply enough), it is the frequency of the driving mechanism, not the amplitude, that may have a more pronounced effect on driving CSF. As Williams^39^ pointed out cardiac pulsation transmits energy to the CSF, while wave propagation depends on pressure-induced differences in motion. Because venous blood and CSF are in equilibrium across venous membranes, venous changes create larger changes in CSF compared to arterial changes^74^. This could be the reason why Takizawa et al. ^37^ observed greater cardiac- than respiratory-driven CSF velocities, and greater respiratory- than cardiac-driven CSF displacement.

Our study participants were naive to mind-body approaches, including breath awareness and breath training. During baseline data collection, most, if not all, of our study participants indicated that they were not aware of any of the different deep breathing practices in our MRI protocol. Thus, the respiratory dynamics investigated in this baseline dataset provides only the immediate influence of yogic breathing in non-practitioners. In our ongoing interventional RCT study, we hypothesized that respiratory dynamics would be different in advanced practitioners, resulting in larger respiratory dynamics, and thus larger effects on CSF. In the RCT study, we will compare pre- and post-intervention respiratory dynamics utilizing the parameters as described herein to determine whether cardiac pulsation is still the primary driver of CSF post-intervention.

Differences in CSF dynamics between individuals, and across breathing conditions within individuals, suggest unique bio-individual characteristics of pulsatile CSF dynamics. Previous studies suggested changes in CSF between individuals could be due to age and sex ^75,76^, vascularization^77^, coupling between arterial inflow and venous outflow^78^. Based on our tests for associations between covariates (age, sex, and BMI), and changes in CSF metrics during SponB versus yogic breathing, we did not observe any significant change except a positive correlation between age and changes in SponB to DCB, in addition to a positive association with age and DCB condition alone. Due to small sample size in our study, we suspect these may be spurious findings. Future studies with larger sample size are needed to explore the associations for these covariates.

Taken together, we demonstrated that pulsatile CSF dynamics are highly sensitive and synchronous to respiratory characteristics such as rate, depth and location of respiratory movement, in agreement with previous studies^13,15,16,37^. Our results provide evidence for immediate modulation of pulsatile CSF dynamics with yogic breathing, and for the importance of studying CSF dynamics in voluntarily controlled conditions to better understand mechanisms driving CSF.

### 4.3. Implications

Understanding the mechanisms that drive CSF dynamics is critical for optimizing brain health and devising potential interventions for disorders of brain waste clearance (e.g., neurodegenerative disorders) as well as for CNS therapeutics (e.g., intrathecal (IT) drug delivery).

Movement of fluids in the CNS have recently drawn major attention for their critical role in the removal of solutes and waste products from the brain interstitium^1,4,7,9,79^. The two discrete fluid compartments of the brain, CSF and ISF, are integral players for CNS homeostasis. While there is a debate whether solutes are removed from the brain through convective flow (suggested by “glymphatic system”^1,18^) or diffusion^19^, CSF-ISF exchange during sleep aids in removal of solutes including amyloid beta^20,21^, a protein associated with Alzheimer’s disease^22^. Emerging epidemiological evidence suggests that sleep disruption is associated with dementia, Alzheimer’s diagnosis ^80,81^ and the development of amyloid plaques prior to the onset of clinical symptoms^82,83^.

Recently, Fultz and colleagues ^9^ conducted neuroimaging in human subjects during sleep by combining blood oxygen level–dependent functional magnetic resonance imaging (BOLD fMRI), electroencephalography (EEG), and CSF flow measurements, and demonstrated (i) CSF flow oscillations during non-rapid eye movement (NREM) sleep were larger and slower (0.05 Hz vasomotion) compared to wakefulness (0.25 Hz respiratory); (ii) slow large CSF waves during NREM sleep were coupled with EEG slow-delta waves and blood oxygenation; (iii) also suggested increased pulsatile CSF dynamics during sleep may alter brain’s waste clearance due to increased mixing and diffusion^2,49^.

In our study, iCSF velocity waveforms were synchronous to breathing patterns, thus slower and larger during slow and deep yogic breathing practices compared to spontaneous breathing. Specifically, we found an increase of 16% - 28% in cranial iCSF velocities during yogic breathing. Because the entry of CSF along perivascular channels is critical for CSF-ISF exchange in rodents^18^, and because increased pulsatile CSF dynamics during sleep potentially alters brain waste clearance due to increased mixing and diffusion, we speculate that increased pulsatile CSF dynamics during yogic breathing, performed during wakefulness, can potentially aid in the removal of waste products in populations regardless of sleep disruption. Validating yogic breathing as a potential therapy for the removal of waste products will then be critical to define the role it may play in the prevention of conditions associated with impaired CSF circulation and/or sleep disruption, such as Alzheimer’s disease.

In addition, increased pulsatile CSF dynamics through yogic breathing could be beneficial for investigating intrathecal (IT) drug delivery and factors influencing IT drug transportation. For instance, using medical image–based computational fluid dynamics Hsu et al. ^84^ studied drug transport as a function of frequency and magnitude of CSF pulsations during different heart rates and CSF stroke volumes. Both heart rate and CSF stroke volume influenced drug distribution in CSF presenting key factors for interpatient variability in drug distribution. We hypothesize that different breathing rates and CSF velocities via yogic breathing would impact peak concentration of drugs in CSF after injection through mixing and diffusion.

### 4.4. Significance

To our knowledge, this is the first report studying modulation of pulsatile CSF dynamics via a mind-body approach^85^. Yogic breathing is a critical component of traditional yoga practices. Yoga has become one of the most popular integrative and complementary mind-body approaches^86^ of the 21^th^ century for cultivating overall health and well-being with an estimated number of 300 million practitioners across the globe^87^. Following the United Nations General Assembly in 2014 ^88^, yoga has been officially recognized as an ‘invaluable gift’ to the world and, to further spread the benefits, June 21 has been formally recognized as International Day of Yoga. In recent decades, yoga has gained the attention of the scientific and clinical communities^60,89^ for its therapeutic benefits, and has become the subject of many research studies for asthma^60^, balance^90^, cognition^91^, stress, anxiety, and depression^56,57^, chronic pain^92,93^, cancer related symptoms ^61,92^, sleep^94^, neurological conditions^95^ such as stroke, multiple sclerosis, epilepsy, traumatic brain injury, Alzheimer’s disease, and more recently management of stress-related problems during the COVID-19 lockdown^96^.

Even though many research studies have demonstrated the positive effects of yoga through specific outcomes^97^, the exact underlying mechanisms for the benefits of yoga are still not fully known. Yoga is a complex multi-modality approach. As often practiced in the West, it is a combination of postures, breathing, relaxation and meditation, making it difficult to determine the specific effects and active mechanisms of isolated yoga practices on specific health outcomes. With an increasing number of practitioners and potentially being suggested as a treatment for several conditions^98^, it is essential to investigate the underlying mechanisms of the active components of yoga to design evidence-based, safe, and targeted practices for specific clinical populations. Nevertheless, yogic breathing, for instance, has been investigated in isolation, and is known to be effective for reducing stress and anxiety^56–58^, lowering blood pressure^59^, improving asthma conditions^60^, and improving response to cancer^61^. However, the impact of yogic breathing in the context of CSF dynamics has never been reported before. Our study may shed light on other components of yoga^97^, and other mind-body approaches with breathing awareness and/or training, such as mindfulness meditation^99^, MBSR^100^, Tai-Chi^101^ or Qi-Gong^102^ in the context of CSF dynamics.

### 4.5. Limitations and Future Studies

Our study is limited by the small sample size. A larger sample size would allow results to be more generalized across all five breathing patterns. The inherent challenges in MRI acquisition can lead to artifacts in MR images. Measurements with data artifacts were removed from the final analysis. Increased temporal resolution in our RT-PCMRI approach results in reduced spatial resolution, which we believe is handled through rigorous data processing methodology including semi-automated algorithm for extracting CSF signals, visually confirming CSF region of interest, and selecting a single voxel. This approach eliminated partial volume effects, but limited CSF velocities within one voxel instead of entire cross section of CSF, and also increased computational cost. Therefore, future work to develop high spatial and temporal resolution for continuous CSF measurements with analysis within the entire CSF region is needed. Despite these limitations, we have shown that our technique can detect and quantify CSF velocities around the spinal cord. To capture true temporal peak velocities and reduce noise due to transient events, we computed averaged peak velocities, which increased computational cost. We have collected pulse data with a finger pulse sensor, and respiration data with a respiration bellow. Future experimental methodology will include electrocardiography (ECG) measures for investigating the heart rate variability (HRV), and potential pressure sensors for measuring intrathoracic and abdominal pressures during yogic breathing techniques. Since our RCT focused on CSF dynamics, we have not investigated arterial and/or venous flow in this study.

In short, future investigations involving yogic breathing and/or other mind-body approaches will need to evaluate the effects of training of the breathing techniques on CSF measures in a formal RCT. Investigations should: (i) use a larger sample size, (ii) study differences in age, sex, gender, race, activity levels, sleep quality, (iii) evaluate the influence of these covariates on pulsatile CSF magnitude and directionality along the spine and in the cranial cavity, (iv) study the coupling between CSF, arterial and venous flow, (v) utilize ECG, intrathoracic and abdominal pressure measurements in sync with MRI, and (vi) evaluate the effect of breathing induced changes in CSF on the brain’s waste clearance mechanism.

## 5. Conclusions

To our knowledge, our study is the first report demonstrating the impact of a mind-body approach such as yogic breathing to modulate CSF dynamics, and comparing with spontaneous breathing. We investigated pulsatile CSF velocities during spontaneous versus yogic breathing practices (slow, deep abdominal, deep diaphragmatic and deep chest breathing) at the level of foramen magnum using a non-invasive MRI-based quantification in a set of healthy participants without current or previous regular practice of mind-body approaches. With rigorous testing, we demonstrated that the three yogic breathing patterns (slow, deep abdominal and deep diaphragmatic) immediately and increased both cranially directed instantaneous CSF velocities and contribution of respiratory power. We observed most statistically significant effects during deep abdominal breathing. Cardiac pulsation was the primary driver of CSF during all breathing conditions except during deep abdominal breathing when there was a comparable contribution of respiration and cardiac 1^st^ harmonic, which suggests respiration can be the primary driver for pulsatile CSF depending on individual differences in breathing depth and location. Since increased pulsatile CSF dynamics is suggested to increase brain’s waste clearance through increased mixing and diffusion, we hypothesize that a potential underlying mechanism for the benefits of yogic breathing is its impact on CSF dynamics, in turn removal of solutes from the brain. Further studies are required to confirm this hypothesis in healthy young and aged brain, and neurological conditions such as Alzheimer’s disease. Our study will shed light on benefits and mechanisms of other components of yoga, and other mind-body approaches in the context of CSF dynamics, and removal of solutes from the brain

## Acknowledgements

We thank study participants for their participation in the study; OCTRI for critical feedback on research plan design; Julie Goldman for assistance in subject recruitment; OHSU AIRC imaging staff for assistance in MRI data acquisition; Ning Jin for initial prototype pulse sequence for MRI acquisition; Michele Hebert, Dr. Mehrad Nazari, Chuck Linke, Katrina Murphy, Dan Klee, and numerous staff and colleagues for the support of the RCT conduct.

## Author contributions

Selda Yildiz: conceptualization, funding acquisition, project administration, supervision, provision, investigation, methodology, data curation, software, formal data analysis/interpretation, visualization, writing the original draft, review and editing; John Grinstead: methodology, review and editing; Andrea Hildebrand: statistical analysis, review and editing; John N. Oshinksi: methodology, data interpretation, review and editing; William D. Rooney: methodology, data interpretation, review and editing; Miranda M. Lim: methodology, data interpretation, review and editing; Barry Oken: methodology, data revision/interpretation, supervision, review and editing.

