## Supplementary Material for "Immediate Impact of Yogic Breathing on Pulsatile Cerebrospinal Fluid Dynamics"

**Affiliations.**
[1]Department of Neurology, Oregon Health & Science University, Portland, OR, 97239, USA
[2]Siemens Medical Solutions USA, Inc, Portland, OR, 97239, USA
[3]Biostatistics and Design Program, Oregon Health & Science University, Portland, OR, 97239, USA
[4] Radiology & Imaging Sciences and Biomedical Engineering, Emory School of Medicine, Emory University, Atlanta, Georgia, 30322, USA
[5]Advanced Imaging Research Center, Oregon Health & Science University, Portland, OR, 97239, USA
[6]Department of Behavioral Neuroscience; Oregon Health & Science University, Portland, OR, 97239, USA
[7]Knight Cardiovascular Institute, Oregon Health & Science University, Portland, OR, 97239, USA
[8]Department of Biomedical Engineering, Oregon Health & Science University, Portland, OR, 97239, USA
[9]VA Portland Health Care System, Portland, OR, 97239, USA
[10]Department of Medicine, Division of Pulmonary and Critical Care Medicine; Oregon Health & Science University, Portland, OR, 97239, USA
[11]Oregon Institute of Occupational Health Sciences; Oregon Health & Science University, Portland, OR, 97239, USA

**This document includes:**

Supplementary text

Figures S1 to S3

Tables S1 to S5

SI References

### Materials and Methods

| Study Inclusion and Exclusion Criteria | |
|---|---|
| **Inclusion Criteria** | • 18-65 years of age<br>• Able to provide consent to be in the study<br>• Willing and able to participate in the study activities<br>• Able to lay supine<br>• Able to walk 15 feet<br>• Naive to mind-body practices, including breath awareness or training in yoga, meditation, Ta-Chi, Qi-Gong<br>• Must have a compatible electronic device for collection of physiological data via a proprietary app* |
| **Exclusion Criteria** | • Inability to provide informed consent<br>• MRI contraindications such as implanted medical devices or other non-removable metal (besides dental fillings), a larger body habitus (e.g., BMI > 30 or weight >250 pounds), claustrophobia<br>• Need for muscle relaxants or anti-anxiety drugs in order to tolerate MRI<br>• Any muscle or skeletal issues that would make it difficult to lie still in the supine position for the 1-hour MRI scan<br>• History of neurological disorders such as stroke, brain hemorrhage, head trauma with loss of consciousness, multiple sclerosis, Parkinson's disease, Huntington's disease, Alzheimer's disease<br>• Sleep disorders<br>• Allergic or respiratory disorders<br>• Craniospinal disorders, e.g., Chiari Malformation.<br>• Spinal injury<br>• Major or uncontrolled psychiatric illness or major depression.<br>• Any condition requiring the use of medication that acts on the brain such as stimulants, sedatives, antidepressants, antipsychotics, and anti-anxiety or anti-seizure medications<br>• Active infection or uncontrolled significant systemic illnesses<br>• Problems with the heart, circulatory system, or lungs such as asthma, chronic obstructive pulmonary disease, pulmonary edema, hypertension, cardiac arrhythmia, unstable angina pectoris, or symptomatic congestive heart failure<br>• Smoking more than 5 cigarettes a day or drinking more than 3 alcoholic beverages a day<br>• Current substance abuse, in drug rehabilitation, or regularly using marijuana<br>• Women only: pregnant or nursing<br>• On the day of MRI scan, fever >100.9$^{o}$F |

**Table S1.** Study Inclusion and Exclusion Criteria. *Our study utilized physiological data devices to objectively track participants' home practice during the 8-week interventions. We excluded participants who did not have a compatible electronic device such as smartphone or tablet.

| MRI Breathing Protocol | | | |
|---|---|---|---|
| | **Breathing Pattern** | **Also Known As** | **Performed** |
| **1** | Spontaneous Breathing (SponB) | Natural Breathing, Resting State Breathing | With or without awareness on inhalation and exhalation without forcing to change the duration and/or the depth of the breath. |
| **2** | Slow Breathing (SlowB) | Slow Rhythmic Breathing | By consciously slowing down the breath with or without deepening the breath. Choice of e.g., 3 to 5 counts* for each inhale/exhale |
| **3** | Deep Abdominal Breathing (DAB) | Belly Breathing, Lower Breathing, Part One of Three-Part Breath* | By consciously creating deep breaths in lower torso by expanding the lower abdomen with inhalation, and relaxing back to resting position with exhalation. Choice of e.g., 3 to 5 counts* for each inhale/exhale |
| **4** | Deep Diaphragmatic Breathing (DDB) | Middle Breathing, Part Two of Three-Part Breath | By consciously creating deep breaths in middle torso by expanding lower ribs and diaphragm to the sides with inhalation and relaxing with exhalation. Choice of e.g., 3 to 5 counts* for each inhale/exhale |
| **5** | Deep Chest Breathing (DCB) | Thoracic Breathing, Upper Breathing, Part Three of Three-Part Breath | By consciously creating deep breaths in upper torso performed by expanding the chest and the lungs to the sides, front and back with inhalation, and relaxing back to resting position with exhalation. Choice of e.g., 3 to 5 counts* for each inhale/exhale |

**Table S2.** MRI Breathing Protocol. *Three-Part Breath. Of the five breathing techniques in the MRI breathing protocol, the last three deep breathing techniques (#3-5) together forms a specific yogic breathing called "*three-part breath*[1]" (Dirgha Pranayama). Typically, these three deep breathing practices are first performed in isolation for a few minutes for practitioners to gain ability to isolate the breathing and associated movement in lower, middle, and upper torso separately, with a long-term goal to improve the breathing apparatus and utilize the full capacity of the lungs. After several rounds of stand-alone performance in each part, practitioners then perform the *three-part breath* by combining all three parts during one breath: "During inhalation, beginning with expanding lower abdomen, smoothly moving the breath and expansion up to diaphragm, then to chest (all in one inhale); and during exhalation, beginning with contracting chest, smoothly moving the breath and contraction down to diaphragm, then to lower abdomen (all in one exhale)”, forming one repetition of a *three-part breath.*

**Real Time - Phase Contrast MRI (RT-PCMRI) CSF voxel selection**

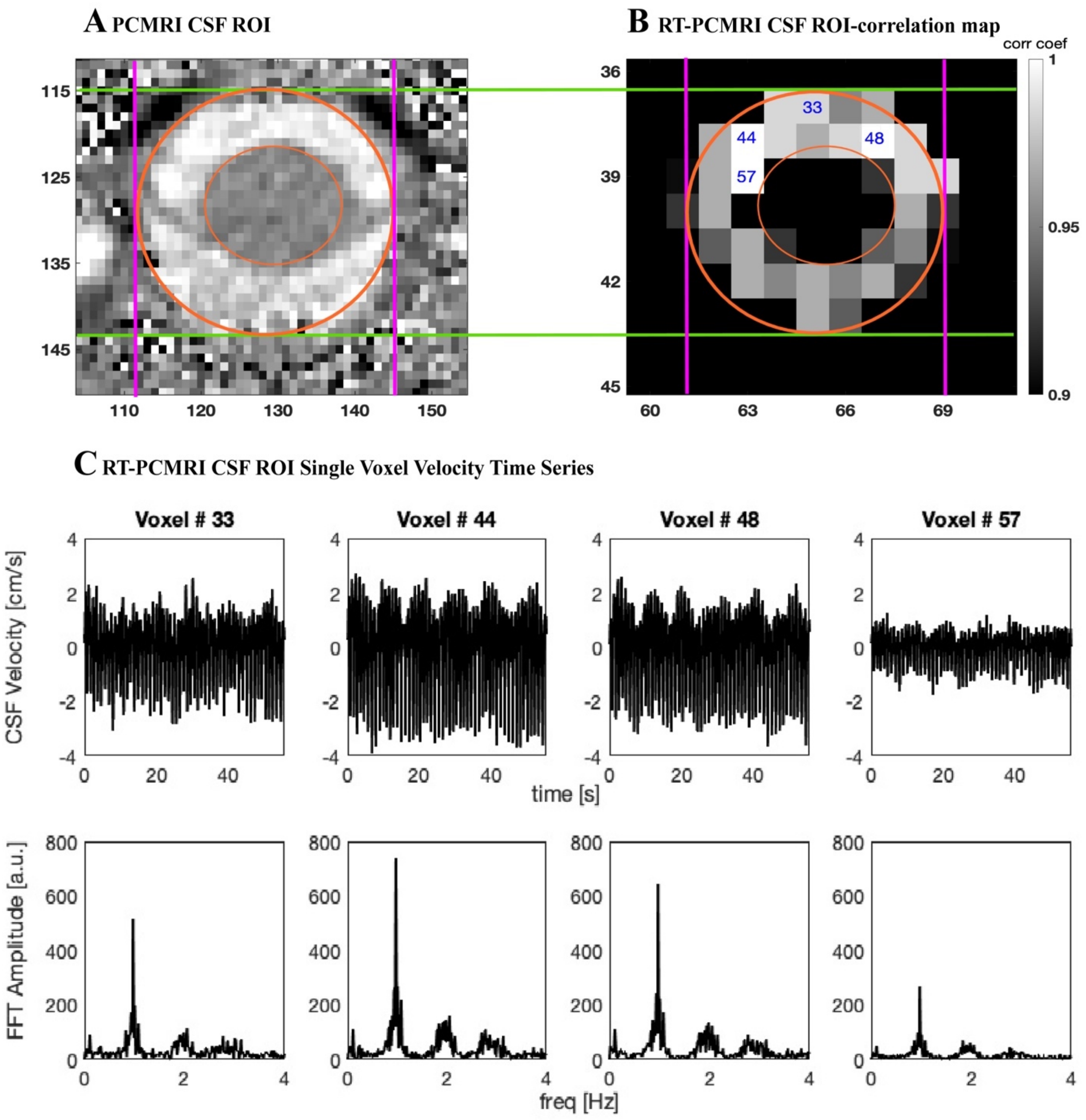


**Figure S1. A**. Sample conventional cardiac-gated PCMRI image (spatial resolution of 0.625 mm x0.625 mm) showing the CSF region of interest (ROI) within orange circles. **B.** Increasing temporal resolution for RT-PCMRI reduces the spatial resolution (2.5 mm x 2.5 mm). To capture true spatial velocity peak values, we implemented a 2-step process. Utilizing our previously developed correlation mapping technique, we first computed highly correlated CSF ROI voxels (greater than 0.7 correlation coefficient) – e.g., 33, 44, 48, 57 as labeled within each ROI. We then compared PCMRI image (A) with RT-PCMRI correlation map (B) to visually confirm the location of voxels. We are interested in maximum capacity of breathing impact on CSF. Anterior CSF velocities were usually greater than posterior velocities across our study population. Therefore, we have selected a single anterior voxel, within the CSF space and not contaminated by partial volume effects, and with greater velocity, which was usually among the highest correlated voxels obtained from the correlation mapping technique (greater than 0.9 correlation coefficient) such as voxel#44 for this sample. **C.** For instance, while voxels #33, 48, and 57 are highly correlated one another, partial volume effects due to spinal cord or outside the subarachnoid space, the velocity waveforms and/or peak values are not truly preserved compared to voxel #44 which is within the CSF space.

### Confirming static tissue voxels are not impacted by deep breathing

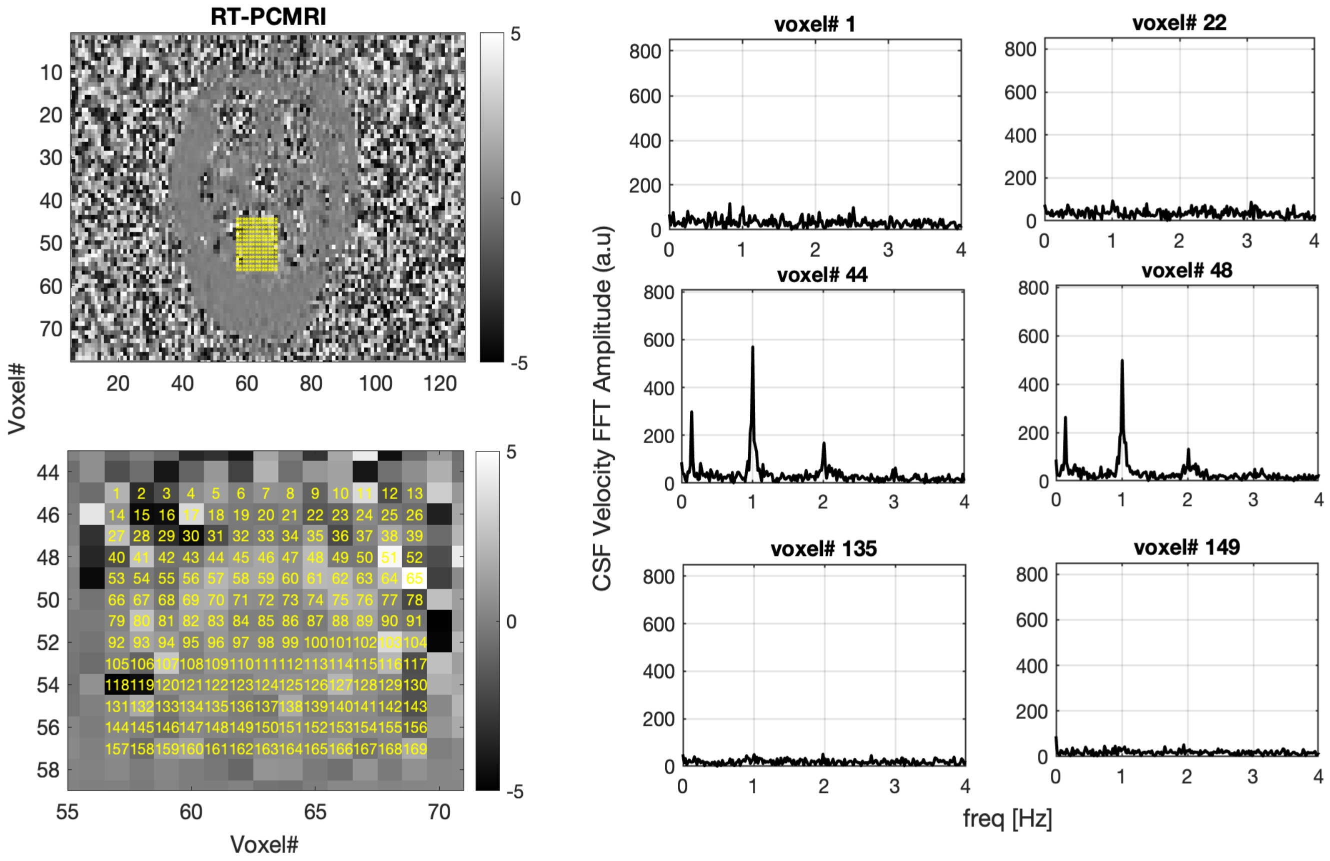


**Figure S2**. To confirm deep breathing practices did not cause any artifacts in CSF velocities, we computed frequency domain signals within static tissue versus CSF region of interest. While we observed peak amplitudes for respiratory and cardiac components in pixels with CSF, we did not observe any respiratory or cardiac components in static tissue.

### Observing cardiac harmonics in frequency domain CSF velocity signals

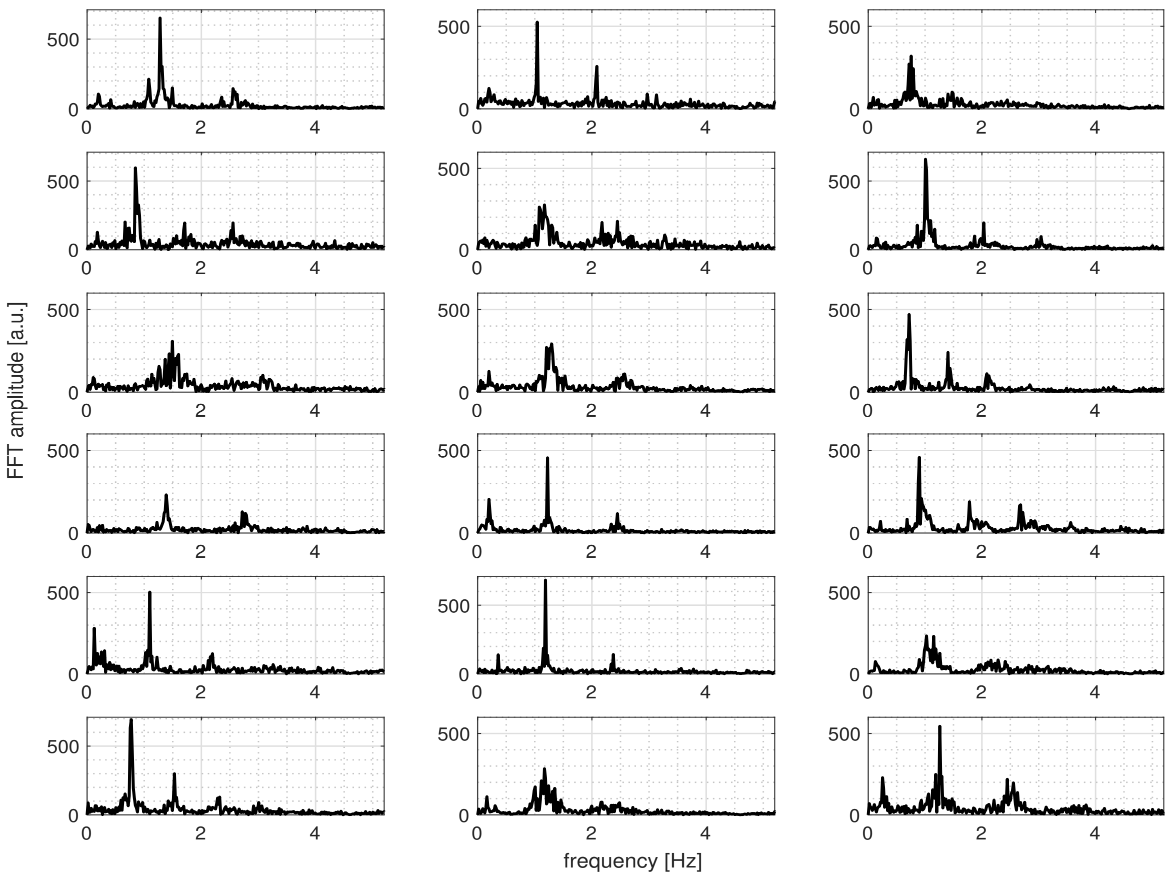


**Figure S3.** Frequency components of CSF Velocities during spontaneous breathing (SponB) for the 18 participants. Previous studies reported vasomotion, respiration, and cardiac (1st harmonic) components of CSF signals, but not cardiac 2nd harmonic component. We observed higher order harmonics of cardiac pulsations (f > ~1.6 Hz) in our preliminary analysis of frequency domain CSF velocity signals. Having observed 1st and 2nd cardiac harmonics but not 3rd or 4th in all subjects, we have included 2nd cardiac harmonics in our analysis because it provides critical information for determining the relative contribution of the underlying mechanisms regulating pulsatile CSF velocities.

### Results

| Time Domain CSF Metrics Assessed for All Breathing Conditions | | | | | | |
|---|---|---|---|---|---|---|
| N=18 | | SponB | SlowB | DAB | DDB | DCB |
| **iCSF** (0-4 Hz) | APMax [cm/s] | 2.10 (0.42) | 2.52 (0.53) | 2.65 (0.56) | 2.57 (0.60) | 2.39 (0.54) |
| | *SponB %Δ* | | 21.55 (23.03) | 28.42 (28.74) | 23.33 (24.49) | 15.57 (26.22) |
| | APMin [cm/s] | -2.74 (0.58) | -3.03 (0.64) | -2.98 (0.69) | -2.87 (0.68) | -2.67 (0.57) |
| | *SponB %Δ* | | 11.28 (15.20) | 9.51 (16.72) | 5.60 (18.46) | -1.94 (10.43) |
| | APMaxMin [cm/s] | 4.84 (0.92) | 5.54 (1.12) | 5.63 (1.11) | 5.44 (1.22) | 5.06 (1.03) |
| | *SponB %Δ* | | 15.34 (16.93) | 17.38 (18.21) | 12.69 (17.36) | 5.23 (14.78) |
| | Disp. [mm] | 0.041 (0.40) | -0.178 (0.35) | -0.146 (0.34) | -0.023 (0.41) | 0.091 (0.43) |
| **rCSF** (0~0.6 Hz) | APMax [cm/s] | 0.68 (0.34) | 0.91 (0.45) | 1.29 (0.54) | 1.14 (0.58) | 1.00 (0.61) |
| | *SponB %Δ* | | 60.27 (90.02) | 118.12 (113.24) | 83.68 (82.02) | 62.92 (90.67) |
| | APMin [cm/s] | -0.67 (0.27) | -0.92 (0.5) | -1.07 (0.42) | -1.05 (0.46) | -0.87 (0.34) |
| | *SponB %Δ* | | 52.90 (88.70) | 77.94 (81.30) | 68.14 (68.01) | 42.57 (62.35) |
| | APMaxMin [cm/s] | 1.36 (0.57) | 1.82 (0.91) | 2.36 (0.88) | 2.19 (0.96) | 1.87 (0.89) |
| | *SponB %Δ* | | 53 (85) | 96 (93) | 73 (72) | 50 (69) |
| | Disp. (mm) | 0.014 (0.22) | -0.108 (0.29) | -0.056 (0.31) | -0.001 (0.24) | 0.073 (0.30) |
| **c1CSF** (0.6~1.7 Hz) | APMax [cm/s] | 1.75 (0.40) | 1.92 (0.45) | 1.80 (0.46) | 1.79 (0.45) | 1.67 (0.39) |
| | *SponB %Δ* | | 10.85 (16.33) | 3.18 (15.5) | 3.69 (18.78) | -3.72 (14.06) |
| | APMin [cm/s] | -1.77 (0.41) | -1.95 (0.48) | -1.81 (0.47) | -1.78 (0.46) | -1.69 (0.41) |
| | *SponB %Δ* | | 10.85 (15.72) | 3.02 (15.79) | 1.72 (16.89) | -4.00 (13.14) |
| | APMaxMin [cm/s] | 3.51 (0.81) | 3.86 (0.93) | 3.61 (0.93) | 3.57 (0.91) | 3.36 (0.80) |
| | *SponB %Δ* | | 10.84 (15.94) | 3.08 (15.54) | 2.68 (17.76) | -3.88 (13.50) |
| | Disp. [mm] | 0.019 (0.08) | 0.007 (0.06 | 0.004 (0.07) | 0.020 (0.09) | 0.025 (0.07) |
| **c2CSF** (0.6~1.7 Hz) | APMax [cm/s] | 0.87 (0.21) | 0.93 (0.23) | 0.95 (0.28) | 0.95 (0.31) | 0.86 (0.22) |
| | *SponB %Δ* | | 7.78 (19.31) | 9.71 (25.94) | 9.47 (27.10) | -0.09 (18.54) |
| | APMin [cm/s] | -0.87 (0.21) | -0.93 (0.23) | -0.95 (0.29) | -0.95 (0.31) | -0.87 (0.23) |
| | *SponB %Δ* | | 7.80 (19.37) | 9.63 (25.97) | 9.31 (27.21) | 0.02 (18.94) |
| | APMaxMin [cm/s] | 1.75 (0.41) | 1.86 (0.46) | 1.91 (0.57) | 1.90 (0.61) | 1.73 (0.46) |
| | *SponB %Δ* | | 7.80 (19.50) | 9.67 (25.93) | 9.39 (27.14) | -0.04 (18.72) |
| | Disp. [mm] | -0.014 (0.03) | 0.004 (0.03) | -0.004 (0.02) | -0.006 (0.02) | -0.008 (0.02) |

**Table S3.** Time Domain CSF metrics (mean, SD, and %Δ compared to SponB). SponB: Spontaneous breathing, SlowB: Slow breathing, DAB: deep abdominal breathing, DDB: deep diaphragmatic breathing, DCB: deep chest breathing, iCSF: instantaneous-CSF, rCSF: respiratory-CSF, $c_1$CSF: 1st cardiac harmonic component of CSF, $c_2$CSF: 2nd cardiac harmonic component of CSF, APMax: averaged peak maximum, APMin: averaged peak minimum, APMaxMin: averaged peak maximum to averaged peak minimum, Disp: displacement. All metrics -except APMaxMin for rCSF, $c_2$CSF, and $c_2$CSF- are used for statistical testing for the differences between SponB and four yogic breathing techniques.

Note that traditional method for computing cranially- and caudally-directed velocities are to compute peak maximum and peak minimum values. In addition to peak maximum and minimum (presented in Table S5), we computed averaged peak maximum and averaged minimum values. Since our goal during each breathing condition is to capture true maximum capacity of CSF velocity, the averaged peak values allowed us to reduce temporal noise caused by i) random transient events that are not part of the regular breathing pattern (e.g., unexpected deep sigh) resulting in greater peak values, or ii) “participant fatigue” experienced during performing slow and/or deep breathing conditions resulting in lower peak values.

| Frequency Domain CSF Metrics Assessed for All Breathing Conditions | | | | | | |
|---|---|---|---|---|---|---|
| **N=18** | | **SponB** | **SlowB** | **DAB** | **DDB** | **DCB** |
| Peak Freq. [Hz] | $rCSF_f$ | 0.18 (0.07) | 0.10 (0.02) | 0.13 (0.04) | 0.13 (0.05) | 0.14 (0.05) |
| | *SponB %Δ* | | -42.44 (18.46) | -23.70 (31.22) | -21.76 (29.69) | -18.72 (33.10) |
| | $c_1CSF_f$ | 1.09 (0.22) | 1.06 (0.25) | 1.12 (0.27) | 1.11 (0.28) | 1.12 (0.26) |
| | *SponB %Δ* | | -2.11 (8.68) | 2.78 (8.58) | 1.78 (10.31) | 2.62 (8.70) |
| | $c_2CSF_f$ | 2.13 (0.40) | 2.09 (0.38) | 2.20 (0.46) | 2.24 (0.49) | 2.26 (0.48) |
| | *SponB %Δ* | | -1.60 (6.02) | 3.21 (8.30) | 4.85 (7.56) | 5.60 (8.77) |
| Peak Amp. [a.u.] | $rCSF_{peak}$ | 114.65 ± 64.98 | 233.61 (177.10) | 302.81 (145.70) | 233.33 (122.37) | 201.74 (135.02) |
| | *SponB %Δ* | | 140.62 (203.42) | 233.59 (244.68) | 160.39 (211.50) | 100.87 (140.32) |
| | $c_1CSF_{peak}$ | 454.71 (162.77) | 403.77 (192.40) | 366.60 (182.68) | 407.04 (204.77) | 390.35 (208.72) |
| | *SponB %Δ* | | -8.63 (33.31) | -19.70 (24.41) | -6.43 (21) | -10.42 (38.55) |
| | $c_2CSF_{peak}$ | 159.42 (64.90) | 123.41 (50.34) | 128.02 (52.32) | 120.78 (52.58) | 130.68 (62.64) |
| | *SponB %Δ* | | -20.39 (19.19) | -15.21 (32.40) | -20.62 (28.54) | -12.86 (36.75) |
| Peak Amp. Ratio | $r/c_{1peak}$ | 0.27 (0.14) | 0.75 (0.83) | 1.16 (1.09) | 0.72 (0.64) | 0.69 (0.75) |
| | *SponB %Δ* | | 193.21 (289.50) | 359.11 (394.92) | 191.82 (200.21) | 158.29 (176.33) |
| | $r/c_{2peak}$ | 0.84 (0.56) | 2.44 (3.06) | 2.95 (2.43) | 2.36 (2.07) | 1.96 (1.89) |
| | *SponB %Δ* | | 223.12 (288.82) | 349.86 (412.26) | 265.23 (332.73) | 165.76 (185.06) |
| Freq. Band [Hz] | $rCSF_{fband}$ | f < 0.60 (0.13) | f < 0.55 (0.10) | f < 0.60 (0.13) | f < 0.60 (0.15) | f < 0.63 (0.16) |
| | $c_1CSF_{fband}$ | 0.60 (0.13) < f < 1.65 (0.33) | 0.55 (0.10) < f < 1.63 (0.30) | 0.60 (0.13) < f < 1.66 (0.33) | 0.60 (0.15) < f < 1.69 (0.35) | 0.63 (0.16) < f < 1.70 (0.34) |
| | $c_2CSF_{fband}$ | 1.65 (0.33) < f < 2.70 (0.56) | 1.63 (0.30) < f < 2.65 (0.56) | 1.66 (0.33) < f < 2.82 (0.59) | 1.69 (0.35) < f < 2.78 (0.60) | 1.70 (0.34) < f < 2.70 (0.58) |
| Power [a.u.] | $rCSF_{power}$ | 1075.26 (975.60) | 2500.00 (3457.83) | 4167.78 (4351.75) | 3141.90 (3054.91) | 2468.69 (2811.90) |
| | *SponB %Δ* | | 229.62 (395.49) | 471.51 (562.65) | 269.51 (314.29) | 187.36 (241.35) |
| | $c_1CSF_{power}$ | 10843.7 (5369.9) | 12442.7 (6417.3) | 10689.4 (5713.3) | 10660.3 (6599.1) | 9440.3 (4700.4) |
| | *SponB %Δ* | | -0.63 (7.59) | 1.34 (9.60) | 2.96 (13.29) | 3.72 (10.83) |
| | $c_2CSF_{power}$ | 2224.13 (1159.82) | 2449.03 (1279.45) | 2407.07 (1379.31) | 2440.93 (1675.36) | 2082.70 (1176.44) |
| | *SponB %Δ* | | 14.88 (35.51) | 11.27 (47.05) | 9.62 (48.13) | -4.19 (34.42) |
| Power Ratio | $r/c_{1power}$ | 0.13 ± 0.15 | 0.29 (0.51) | 0.59 (0.78) | 0.43 (0.56) | 0.40 (0.60) |
| | *SponB %Δ* | | 247.46 (617.84) | 533.68 (866.40) | 294.27 (348.82) | 249.35 (299.46) |
| | $r/c_{2power}$ | 0.63 ± 0.81 | 1.72 (3.52) | 2.85 (4.38) | 2.16 (2.96) | 1.75 (2.39) |
| | *SponB %Δ* | | 324.26 (974.70) | 589.40 (1153) | 327.78 (513.69) | 234.62 (319.57) |

**Table S4.** Frequency Domain CSF metrics (mean, SD, and %Δ compared to SponB). SponB: spontaneous breathing, SlowB: slow breathing, DAB: seep abdominal breathing, DDB: deep diaphragmatic breathing, DCB: deep chest breathing, rCSF: respiratory-CSF, $c_1$CSF: 1st cardiac harmonic component of CSF, $c_2$CSF: 2nd cardiac harmonic component of CSF, APMax: averaged peak maximum, APMin: averaged peak minimum, APMaxMin: averaged peak maximum to averaged peak minimum, Disp: displacement, $rCSF_f$: peak frequency of rCSF, $c_1CSF_f$: peak frequency of $c_1$CSF, $c_2CSF_f$: peak frequency of $c_2$CSF, $rCSF_{peak}$: peak frequency amplitude of rCSF, $c_1CSF_{peak}$: peak frequency amplitude of $c_1$CSF, $c_2CSF_{peak}$: peak frequency amplitude of $c_2$CSF, $r/c_{1peak}$: peak frequency amplitude ratio of rCSF to $c_1$CSF, $r/c_{2peak}$: peak frequency amplitude ratio of rCSF to $c_2$CSF, $rCSF_{power}$: power of rCSF, $c_1CSF_{power}$: power of $c_1$CSF, $c_2CSF_{power}$: power of $c_2$CSF, $r/c_{1power}$: power ratio of rCSF to $c_1$CSF, $r/c_{2power}$: power ratio of rCSF to $c_2$CSF. All metrics -except frequency band- are used for statistical testing for the differences between SponB and four yogic breathing techniques.

| Time Domain: Peak Maximum, Peak Minimum, and Peak Maximum to Peak Minimum Assessed for All Breathing Conditions | | | | | | |
|---|---|---|---|---|---|---|
| | N=18 | SponB | SlowB | DAB | DDB | DCB |
| **iCSF** (0-4 Hz) | PMax [cm/s] | 2.61 (0.53) | 3.06 (0.69) | 3.23 (0.70) | 3.11 (0.65) | 2.90 (0.61) |
| | *SponB %Δ* | | 19.09 (23.11) | 27.12 (30.41) | 21.35 (23.97) | 14.26 (28.37) |
| | PMin [cm/s] | -3.23 (0.62) | -3.63 (0.78) | -3.55 (0.81) | -3.50 (0.82) | -3.23 (0.63) |
| | *SponB %Δ* | | 13.12 (19.12) | 10.56 (20.33) | 9.68 (24.94) | 0.70 (12.06) |
| | PMaxMin [cm/s] | 5.84 (1.08) | 6.69 (1.41) | 6.78 (1.30) | 6.61 (1.37) | 6.12 (1.09) |
| | *%Δ (SponB)* | | 15.30 (18.40) | 17.76 (21.69) | 14.43 (21.47) | 6.33 (17.05) |
| **rCSF** (0~0.6 Hz) | PMax [cm/s] | 0.87 (0.45) | 1.12 (0.50) | 1.49 (0.57) | 1.26 (0.65) | 1.10 (0.66) |
| | *SponB %Δ* | | 59.88 (91.29) | 99.58 (97.06) | 64.40 (76.88) | 47.73 (91.27) |
| | PMin [cm/s] | -0.85 (0.35) | -1.13 (0.65) | -1.30 (0.44) | -1.22 (0.42) | -1.11 (0.47) |
| | *SponB %Δ* | | 65.79 (157.23) | 83.03 (116.76) | 64.54 (76.13) | 58.73 (107.36) |
| | PMaxMin [cm/s] | 1.72 (0.70) | 2.25 (0.10) | 2.78 (0.82) | 2.48 (0.99) | 2.21 (0.98) |
| | %Δ (SponB) | | 54.48 (108.75) | 87.26 (100.41) | 58.94 (70.53) | 46.55 (85.88) |
| **c1CSF** (0.6 ~1.7 Hz) | PMax [cm/s] | 2.02 (0.46) | 2.21 (0.52) | 2.09 (0.51) | 2.13 (0.51) | 1.97 (0.43) |
| | *SponB %Δ* | | 11.25 (19.33) | 4.65 (19.84) | 7.97 (28.09) | -0. 89 (17.31) |
| | PMin [cm/s] | -2.00 (0.43) | -2.31 (0.62) | -2.11 ± 0.54 | -2.07 (0.53) | -1.98 (0.46) |
| | *SponB %Δ* | | 15.97 (19.38) | 6.57 (21.15) | 4.99 (24.68) | -0.36 (16.28) |
| | PMaxMin [cm/s] | 4.02 (0.87) | 4.53 (1.13) | 4.20 (1.04) | 4.20 (1.02) | 3.95 (0.88) |
| | *SponB %Δ* | | 13.43 (18.42) | 5.45 (20.03) | 6.32 (25.89) | -0.85 (15.67) |
| **c2CSF** (0.6 ~1.7 Hz) | PMax [cm/s] | 1.06 (0.26) | 1.16 (0.29) | 1.22 (0.36) | 1.21 (0.40) | 1.07 (0.26) |
| | *SponB %Δ* | | 12.05 (25.59) | 16.68 (31.08) | 15.76 (31.79) | 3.16 (23.15) |
| | PMin [cm/s] | -1.05 (0.26) | -1.16 (0.30) | -1.23 (0.36) | -1.24 (0.42) | -1.08 (0.31) |
| | *SponB %Δ* | | 13.28 (23.63) | 18.05 (28.91) | 19.73 (36.70) | 4.68 (25.75) |
| | PMaxMin [cm/s] | 2.11 (0.51) | 2.32 (0.58) | 2.45 (0.72) | 2.45 (0.81) | 2.15 (0.57) |
| | *SponB %Δ* | | 12.55 (24.07) | 17.25 (29.43) | 17.67 (34.00) | 3.83 (23.94) |

**Table S5.** Time domain peak maximum, peak minimum, and peak maximum to peak minimum metrics were calculated in addition to averaged peak CSF metrics (mean, SD, and %Δ compared to SponB). SponB: spontaneous breathing, SlowB: slow breathing, DAB: seep abdominal breathing, DDB: deep diaphragmatic breathing, DCB: deep chest breathing, rCSF: respiratory-CSF, c1CSF: 1st cardiac harmonic component of CSF, c2CSF: 2nd cardiac harmonic component of CSF, APMax: averaged peak maximum, APMin: averaged peak minimum, APMaxMin: averaged peak minimum to averaged peak minimum.
